# Nascent RNA sequencing reveals a dynamic global transcriptional response at genes and enhancers to the natural medicinal compound celastrol

**DOI:** 10.1101/117689

**Authors:** Noah Dukler, Gregory T. Booth, Yi-Fei Huang, Nathaniel Tippens, Charles G. Danko, John T. Lis, Adam Siepel

## Abstract

Most studies of responses to transcriptional stimuli measure changes in cellular mRNA concentrations. By sequencing nascent RNA instead, it is possible to detect changes in transcription in minutes rather than hours, and thereby distinguish primary from secondary responses to regulatory signals. Here, we describe the use of PRO-seq to characterize the immediate transcriptional response in human cells to celastrol, a compound derived from traditional Chinese medicine that has potent anti-inflammatory, tumor-inhibitory and obesity-controlling effects. Our analysis of PRO-seq data for K562 cells reveals dramatic transcriptional effects soon after celastrol treatment at a broad collection of both coding and noncoding transcription units. This transcriptional response occurred in two major waves, one within 10 minutes, and a second 40-60 minutes after treatment. Transcriptional activity was generally repressed by celastrol, but one distinct group of genes, enriched for roles in the heat shock response, displayed strong activation. Using a regression approach, we identified key transcription factors that appear to drive these transcriptional responses, including members of the E2F and RFX families. We also found sequence-based evidence that particular TFs drive the activation of enhancers. We observed increased polymerase pausing at both genes and enhancers, suggesting that pause release may be widely inhibited during the celastrol response. Our study demonstrates that a careful analysis of PRO-seq time course data can disentangle key aspects of a complex transcriptional response, and it provides new insights into the activity of a powerful pharmacological agent.

## Introduction

The technique of perturbing cells and then measuring changes in their patterns of gene expression is a reliable and widely used approach for revealing mechanisms of homeostatic regulation. In mammalian cells, a wide variety of stimuli that induce striking changes in transcription are routinely applied, including heat shock, hormones such as estrogen, androgen and cortisol, lipopolysaccharide, and various drugs. Regardless of the stimulus, transcription is commonly assayed by measuring concentrations of mature mRNA molecules, typically using RNA-seq. This approach is now relatively straightforward and inexpensive, and allows for the use of standard analysis tools in detecting many transcriptional responses^1,2^.

Nevertheless, these mRNA-based approaches are fundamentally limited in temporal resolution owing to the substantial lag between changes in transcriptional activity and detectable changes in the level of mRNAs. This lag results in part from the time required for transcription and post-transcriptional processing, and in part because pre-existing mRNAs buffer changes in mRNA concentration. For a typical mammalian gene, significant changes may require hours to detect, making it difficult to distinguish primary responses to a signal from secondary regulatory events. A possible remedy for this limitation is instead to make use of GRO-seq^3^, PRO-seq^4^, NET-seq^5–7^, or related methods^8–11^ for assaying nascent RNAs. These assays have the important advantage of directly measuring the production of new RNAs, rather than concentrations of mature mRNAs. As a consequence, they can detect immediate changes in transcriptional activity, and they permit time courses with resolutions on the order of minutes rather than hours^12–15^. An additional benefit of nascent RNA sequencing is that it is effective in detecting unstable noncoding RNAs, including enhancer RNAs (eRNAs), together with protein-coding transcription units^13,16,17^. As a result, both active regulatory elements (which are generally well marked by eRNAs) and transcriptional responses can be detected using a single assay^18^.

In this study, we sought to use PRO-seq to characterize the immediate, dynamic transcriptional response to the compound celastrol. Celastrol (also known as tripterine) is a pentacyclic triterpenoid isolated from the root extracts of *Tripterygium wilfordii* (thunder god vine), which has been used for millennia in traditional Chinese medicine for treatment of fever, joint pain, rheumatoid arthritis, bacterial infection, and other ailments^19^. During the past few decades, celastrol has shown promise as an anti-inflammatory agent in animal models of collagen-induced arthritis, Alzheimer disease, asthma, systemic lupus erythematosus, and rheumatoid arthritis^20–24^. In addition, it is known to inhibit the proliferation of tumor cells, including those from leukemia, gliomas, prostate, and head/neck cancer^24–27^. Recent research has also demonstrated striking obesity-controlling effects in mice^28,29^. Celastrol is known to activate the mammalian heat shock transcription factor HSF1 and stimulate the heat shock response^19,30^ as well as the unfolded protein response^27,31^, and to activate a battery of antioxidant response genes^30^. Celastrol also inhibits the activities of other transcription factors, including androgen receptor^32^, glucocorticoid receptor^30^ and NF-KB^24^. Nevertheless, much remains unknown about the mechanism of action by which celastrol affects transcription. In particular, little is known about the immediate transcriptional effects or primary targets of celastrol. Thus, an examination using PRO-seq provides an opportunity for a deeper understanding of the mechanisms underlying the activity of this potent compound which has potential therapeutic implications.

We collected PRO-seq data for K562 cells at tightly spaced time points after treatment with celastrol and analyzed these data using a variety of computational methods. Our analysis revealed dramatic transcriptional effects within minutes of celastrol treatment at a large, diverse collection of both coding and noncoding transcription units. We observed a general pattern of transcriptional repression, but saw activation in a group of genes enriched for roles in the heat shock response. By analyzing the associated sequences, we were able to identify numerous transcription factors that appeared to drive the transcriptional responses at both genes and enhancers. Interestingly, we also observed a clear impact from celastrol treatment on polymerase pausing at both genes and enhancers, suggesting that pause release may be widely inhibited during the celastrol response. Finally, changes in pausing were negatively correlated with changes in transcriptional activity, suggesting that pausing may contribute to the response to celastrol. Altogether, our analysis demonstrates that time-courses of PRO-seq data together with appropriate bioinformatic analyses can be used to dissect key aspects of a complex transcriptional response at both regulatory elements and target genes.

## Results

### Celastrol induces broad transcriptional repression and more limited up-regulation

We prepared PRO-seq libraries for K562 cells before celastrol treatment and after 10, 20, 40, 60, and 160 minutes of celastrol treatment, with two biological replicates per time point (Figure 1A). To ensure that we could normalize read counts even in the presence of global changes in transcription, we spiked in to each sample an equal number of permeable *Drosophila* cells prior to run-on^33^. Samples were sequenced to a total combined depth of 334.3M reads, with an average replicate concordance of *r*^2^≈98% (Supplemental Figure 1). About 0.5M of these reads (0.1%) were derived from the *Drosophila* spike in. To obtain gene models appropriate for our cell types and conditions, we developed a probabilistic method, called tuSelector, that considers all annotated isoforms for each gene and identifies the most likely gene model given our PRO-seq data (Supplemental Figure 2, Supplementary Methods). We identified a total of 12,242 protein-coding genes from GENCODE as being actively transcribed in one or more time-points (Methods). Of these genes, 75.4% were active across all six time points, 11.7% were active in a single time-point, and the remaining 12.9% were active in 2-5 time-points. Thus, our PRO-seq data and computational analyses indicate that more than half of all protein-coding genes are transcribed either in the basal condition or during the celastrol response in K562 cells.

**Figure 1.**
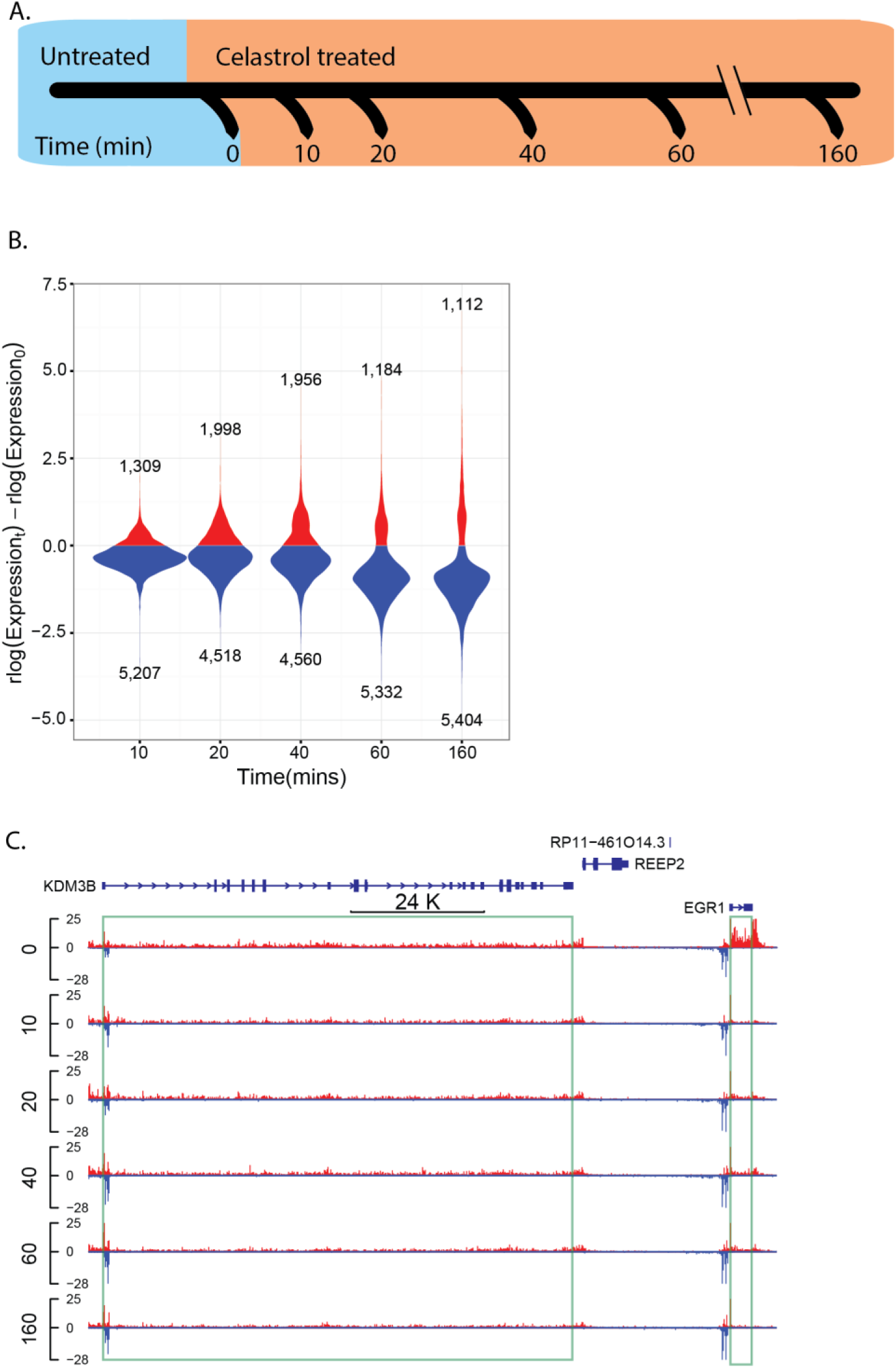
Characterizing the dynamic transcriptional response to celastrol using PRO-seq. **(A)** PRO-seq was applied to K562 cells collected before celastrol treatment (untreated or 0 minutes) and at 10, 20, 40, 60, and 160 minutes after celastrol treatment. Two biological replicates were performed for each time point **(B)** Distribution of log expression ratios (treated vs. untreated) for each time point (rlog is a regularized log_2_ estimate obtained from DESeq2). Only genes classified as differentially expressed (DE) throughout the time course are represented. Notice that most DE genes (FDR ≤ 0.01) are down-regulated upon celastrol treatment. **(C)** A UCSC Genome Browser display showing raw PRO-seq data for two differentially expressed genes, EGR1 and KDM3B. EGR1 is rapidly and strongly repressed (immediate decrease of ~80%), whereas KDM3B is more gradually repressed, losing ~50% of its expression by 160 minutes.

The genes that are differentially transcribed in response to celastrol were of particular interest for further analysis. To measure transcriptional activity specific to each time point, we used counts of PRO-seq reads mapping to the first 16 kb of each gene body. Because RNA polymerase travels at an average rate of ~2 kb/min^12,14,34,35^ and our time points are separated by at least 10 minutes, this strategy conservatively considers new transcription only, yet maintains sufficient statistical power for downstream analysis (see Methods). By applying DEseq2^36^ to these 16-kb read counts, we identified 6,516 (56%) of the active genes as being differentially expressed (DE) relative to the untreated condition (FDR ≤ 0.01). Interestingly, ~80% of these DE genes were down-regulated. Many genes showed rapid and dramatic down-regulation, with decreases in expression by half or more at 3.5% of DE genes within 10 minutes, at 7.8% of DE genes within 20 minutes, and at 48.1% of DE genes within 160 minutes (Figure 1B&C). By contrast, many fewer genes showed substantial increases in expression; for example, only 0.03%, 1.9%, and 7.7% of DE genes had doubled in expression after 10, 20, and 160 minutes, respectively. Nevertheless, extreme up- and down-regulation were both rare, with <1% of DE genes showing increases and <1% showing decreases in transcription by factors of eight or more. These observations are reminiscent of findings for the heat shock response, which have included general decreases in transcription together with up-regulation of selected stress-response elements^15,37^, but the effect of celastrol is somewhat less dramatic. We conclude that celastrol broadly inhibits transcription within minutes after administration but also rapidly activates a set of genes that may be important for continued cellular viability.

### Celastrol activates heat shock more strongly and directly than it activates the unfolded protein response

Celastrol has been reported to activate stress response pathways such as the heat shock and unfolded protein responses^19,27,30,31^. To see whether these effects were detectable at the transcriptional level immediately after treatment with celastrol, we examined our PRO-seq data at genes activated by heat shock factor protein 1 (HSF1) and genes involved in the three branches of the unfolded protein response (UPR), corresponding to activating transcription factor 6 (ATF6), inositol-requiring enzyme 1 (IRE1), and endoplasmic reticulum kinase (PERK) (Figure 2A). Because the initial stages of the UPR and heat shock response are non-transcriptional, we looked for downstream activity of the first group of transcription factors activated in each pathway, using targets reported in the Reactome pathway database^38^. Most direct targets of HSF1 were up-regulated within 160 minutes (Figure 2B). Genes encoding chaperone protein HSP110 and proteinase inhibitor CBP1 were among the HSF1 targets showing the strongest initial response, with the gene encoding HSP110 almost quadrupling its expression in 10 minutes and that for CBP1 increasing more than eight-fold in 160 minutes. Most direct targets of ATF4, ATF6, and XBP1, however, were not strongly induced during our time course. There were some exceptions to this general rule, such as genes encoding transcription factor ATF3, chaperone BiP, and apoptosis inhibitor DNAJB9, which more than doubled in expression. It is possible that these targets are activated earlier than other targets, perhaps by other TFs. In any case, our observations suggest that celastrol induces a pronounced, rapid transcriptional response in the heat shock pathway, and has a much less pronounced transcriptional effect on the UPR, although some targets of the UPR are activated.

**Figure 2.**
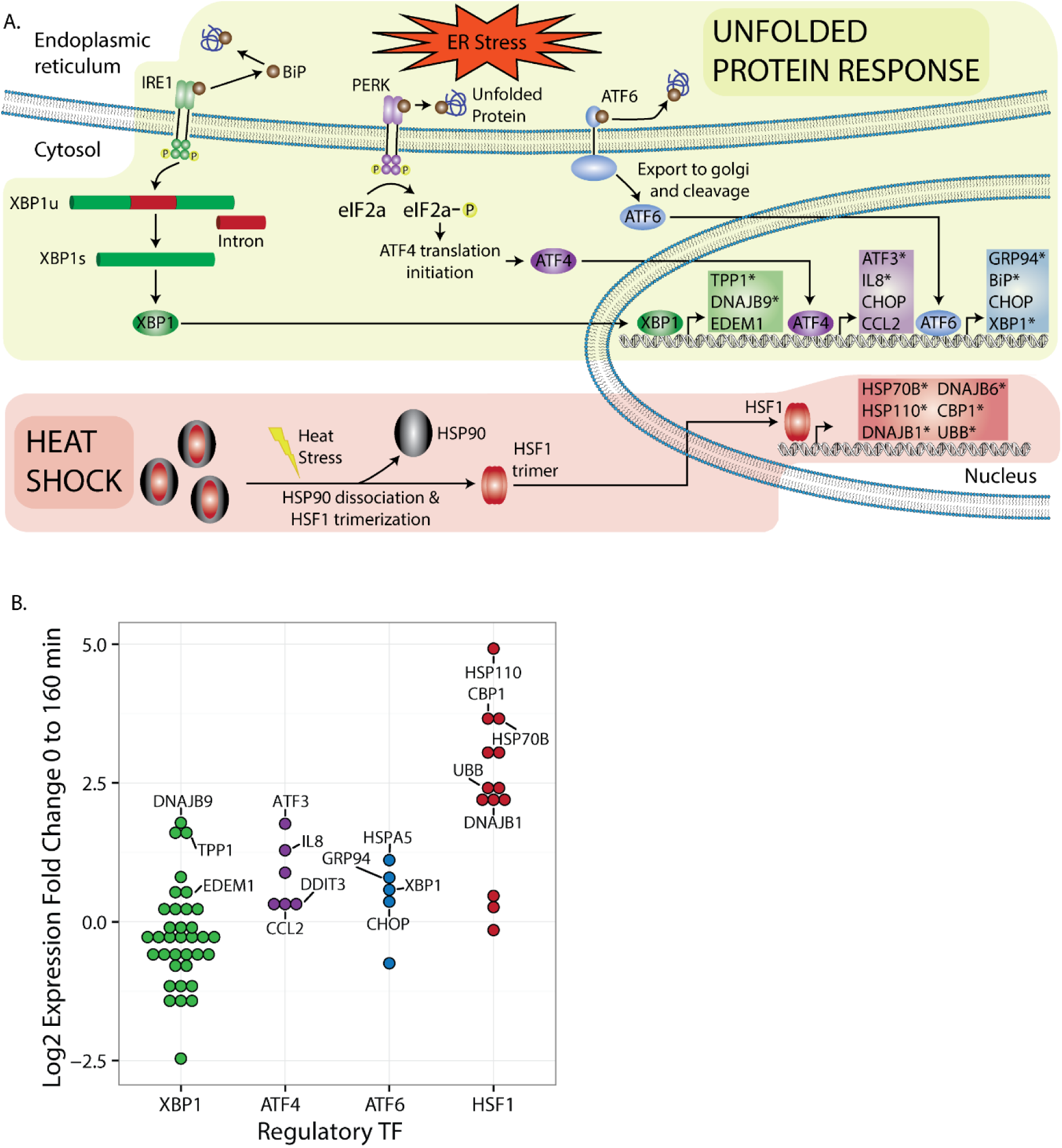
Induction of cellular stress responses by celastrol. **(A)** Illustration showing key aspects of the unfolded protein response (UPR) and heat shock response (HSR), both of which have been reported to be induced by celastrol^30,31^. Expected transcriptional targets are shown inside the nucleus, with targets of HSF1, the key transcription factor (TF) activated in the HSR, in red, and targets of the TFs associated with the three major branches of the UPR—XBP1, ATF4, and ATF6—in green, purple, and blue, respectively. Asterisks indicate genes that were differentially expressed in our experiments with FDR ≤ 0.01. **(B)** PRO-seq-based log fold changes in expression in K562 cells after 160 minutes of treatment by celastrol for numerous known targets of the same four TFs: XBP1, ATF4, ATF6, and HSF1. Only targets of HSF1 display strong up-regulation.

### Celastrol produces distinct temporal patterns of transcriptional response

Our PRO-seq data for closely spaced time points enabled us to examine the temporal patterns of transcriptional response to celastrol treatment across the genome. Using an autoregressive clustering algorithm, EMMIX-WIRE^39^, we partitioned our ~12,000 DE genes into six distinct clusters, based on their patterns of relative transcription across the five time points following celastrol treatment (see Methods) (Figure 3A). Only one of these clusters (cluster #3) displayed dramatic and sustained up-regulation. By contrast, cluster #4 showed transient, weak activation followed by down-regulation, cluster #2 showed rapid and pronounced down-regulation, clusters #1 and #6 displayed moderate, continuous down-regulation, and cluster #5 showed delayed down-regulation. Interestingly, the change points in the expression patterns for these clusters suggested that the transcriptional response to celastrol occurs largely in two distinct waves, one within the first ten minutes, and a second between 40 and 60 minutes. These findings were robust to the number of clusters selected, with similar overall behavior for five-, six-, and seven-cluster models (Supplemental Figure 3&4).

**Figure 3.**
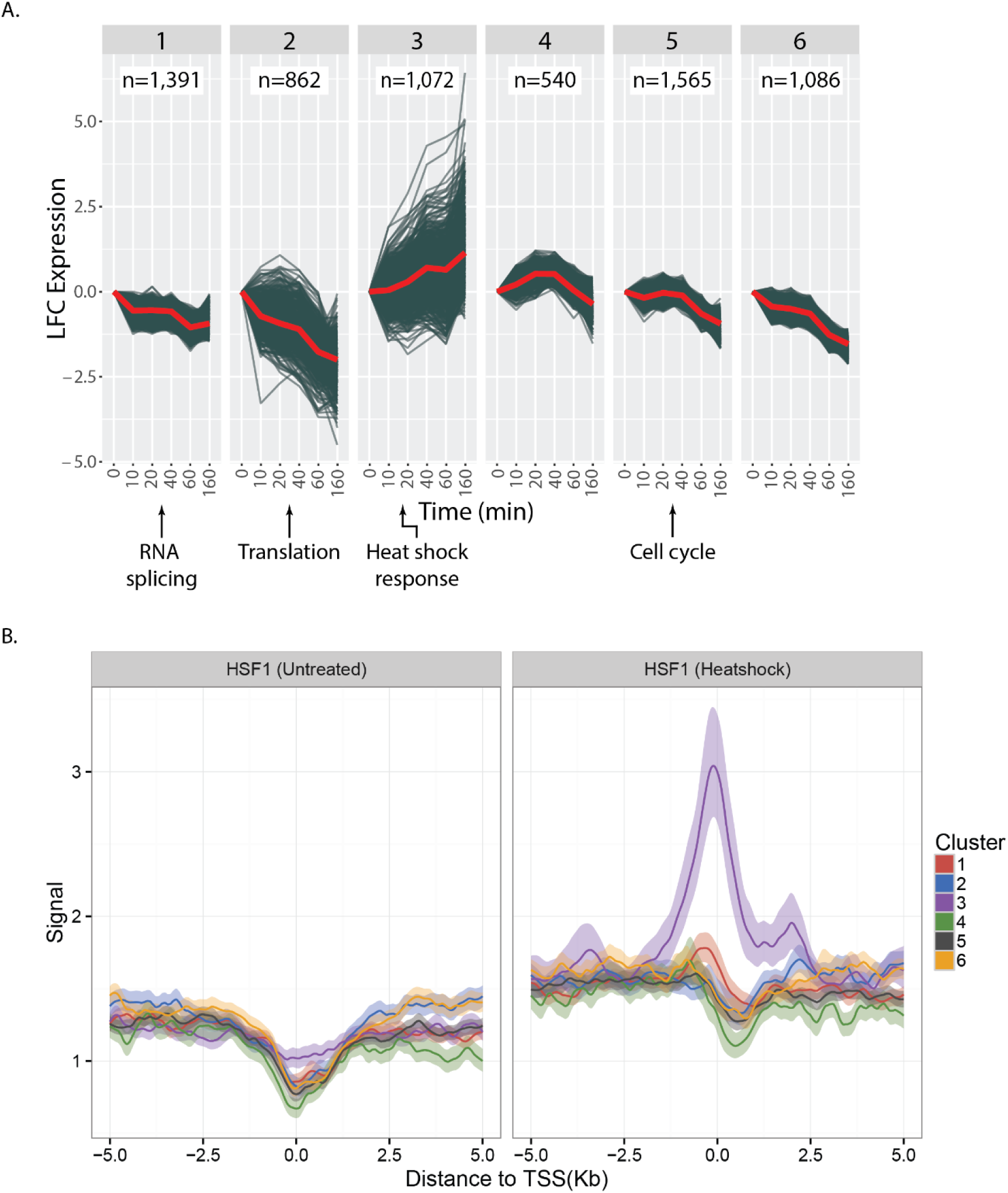
Clusters of genes showing distinct temporal patterns of response to celastrol. **(A)** Differentially expressed genes (FDR ≤ 0.01) clustered by time series of log fold change (LFC) in expression relative to the untreated condition (0 minutes). Each gene is represented by a blue line, and the red lines indicate the mean expression per timepoint per cluster. Below each cluster is a summary of the most strongly enriched terms in the Reactome ontology^38^ (FDR ≤ 0.01; see Supplementary Material for details). Clusters #4 and #6 did not show coherent enrichments. **(B)** ChIP-seq data from reference ^42^ describing binding of HSF1 in K562 cells under normal (left) and heatshock (right) conditions, stratified by our cluster assignments. Each line represents an average over all genes in the cluster in the region of the TSS, with lighter-colored bands representing 95% confidence intervals obtained by bootstrap sampling. Notice that cluster three is unique in showing a strong enrichment for heatshock-induced binding of HSF1.

Four of these six clusters could be associated with fairly coherent biological functions, according to the Reactome ontology (Fig 3A). In particular, cluster #1 contains essential elements of the RNA splicing machinery (e.g., CD2BP2, CLP1, and the SRSF kinase family) (Supplemental Figure 5). Down-regulation of this cluster is consistent with previous reports that splicing is inhibited under heat shock^40^. Cluster #2 is enriched for a wide variety of terms corresponding to ribosomal assembly, translational initiation, and peptide elongation. The pronounced transcriptional repression of this cluster, together with activity of HSPB1 (HSP27) and HSPA2 (HSP70), is consistent with reports that celastrol activates the heat shock response and thereby inhibits translation via HSPB1 in the absence of HSP70^41^ (Supplemental Figure 6&7). Cluster #3 is enriched for genes responsible for the HSF1 response, including the *HSPA* family (Supplemental Figure 8). In addition, many genes in this cluster have been shown, by ChIP-seq, to bind by HSF1 under heat shock conditions in K562 (Figure 3B)^42^. Cluster #4 is enriched for genes associated with cargo transport (Supplemental Figure 9). Finally, cluster #5 is enriched for pathways that enable DNA replication (e.g., MCM family) and cell cycle progression (e.g., CDK family; Supplemental Figure 10). The delayed down-regulation of these genes may occur as the cell is preparing to enter replicative arrest and, potentially, senescence, consistent with observations that celastrol induces cell cycle arrest and potentiates apoptosis^27,43,44^. Clusters #4 and #6 did not exhibit striking functional enrichments. Similar patterns of enrichment were observed with five- and seven-cluster models. Overall, these results demonstrate that the components of a complex, multi-layered transcriptional response can be disentangled, to a degree, by identifying groups of genes that display distinct temporal patterns of gene expression.

### Several key transcription factors are associated with the celastrol response

Can the distinct transcriptional responses in these clusters of genes be traced to particular transcription factors (TFs)? To address this question, we used linear regression to explain the estimated transcription levels at each time point based on the TFs that apparently bind in the promoter region of each gene. We used two orthogonal sources of information about TF binding: (1) ChIP-seq peaks for untreated K562 cells^45^; and (2) sequence-based computational predictions based on DeepBind^46^. In both cases, we considered the interval between 500bp upstream and 200bp downstream of the transcription start site of each active gene. Our regression model included a coefficient for each TF at each time point. A positive estimate of this coefficient indicated that a given TF and its binding sites were associated with up-regulation at a given time point, whereas a negative estimate indicated that the TF and binding sites were associated with down-regulation at that time point.

Between the two TF binding datasets, we identified over twenty TFs as being significantly associated with changes in gene expression and having a large change in effect size (Figure 4, Supplemental Figure 11). Of these TFs, E2F4 stood out as showing a particularly pronounced impact on expression in both datasets. E2F4 is associated with incrementally increased expression between 0 and 60 minutes, and with decreased expression thereafter, similar to the expression pattern of genes in cluster five, which are associated with cell cycle control. This observation is consistent with reports that E2F4 is an activator in some contexts but primarily acts as a repressor responsible for maintaining G2 arrest^47,48^.

**Figure 4.**
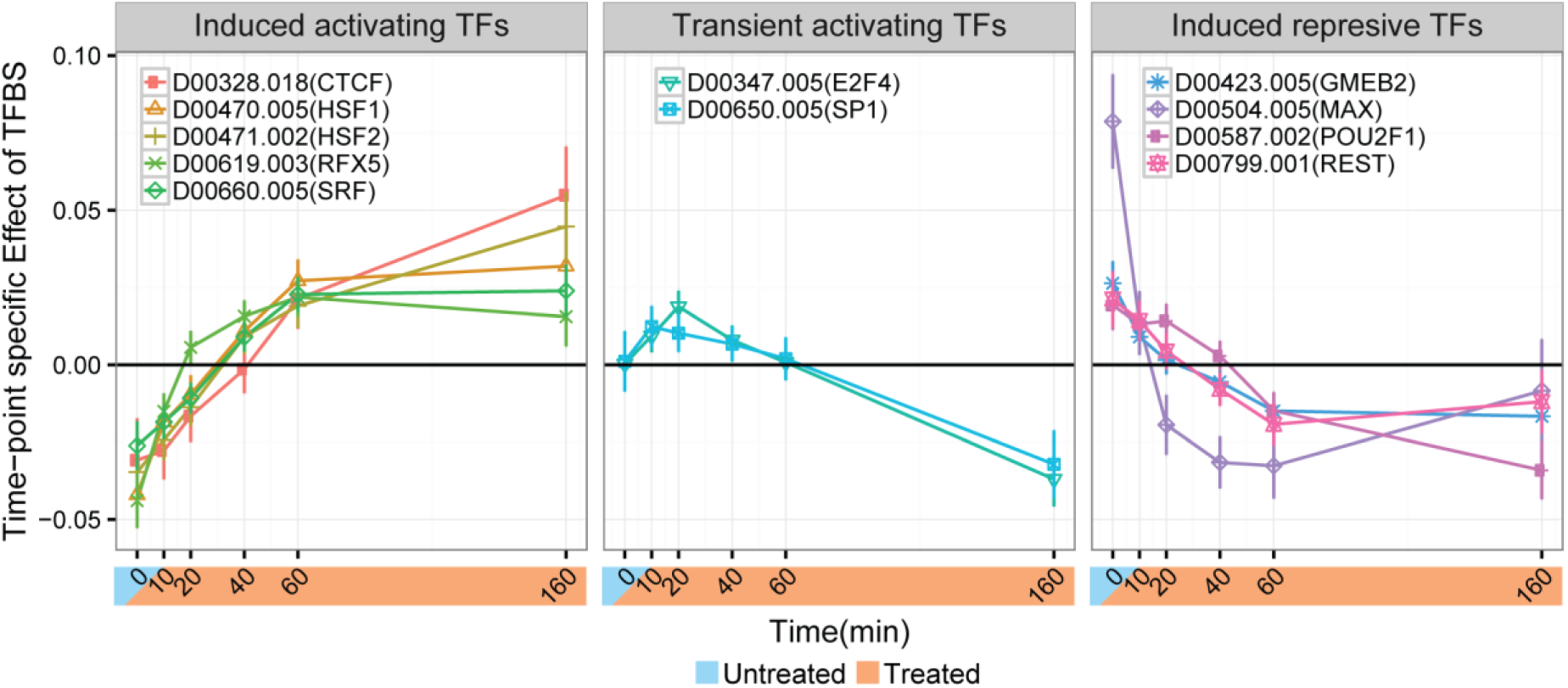
The celastrol response appears to be influenced by various transcription factors (TFs). Shown are the estimated weights from the regression model to predict PRO-seq-measured transcription levels from TF-specific DeepBind scores in the promoter regions of differentially expressed genes. A positive weight for a TF at given timepoint indicates that genes at which that TF is predicted to be bound showed increased expression relative to those without it. Negative weights indicates decreased expression. The three separate plots represent clusters of TFs associated with distinct temporal patterns.

In addition, we found that the dimerizing TFs MYC (from ChIP-seq data) and MAX (from DeepBind predictions) were both associated with an immediate increase in gene expression, followed by decreased expression within 20 minutes (Figure 4). This delayed decrease in expression of MYC- and MAX-bound genes could result from the known disruption of MYC-MAX dimerization by celastrol^19,49^. Finally, genes predicted to be bound by SRF also displayed elevated gene expression after 40 minutes. SRF has been previously associated with early and transient induction of cytoskeletal genes in response to heat stress in murine embryonic fibroblasts^15^.

Finally, we found that the paralogous TFs RFX1 (from ChIP-seq) and RFX5 (from DeepBind) were both associated with increases in expression. Both of these TFs have been implicated in regulating MHC-II expression and both have context-specific transcriptional repression and activation mechanisms^50–52^, so it is possible that they contribute to celastrol’s anti-inflammatory effects. However, RFX1 was not tested with DeepBind and its motif is quite similar to that of RFX5 so it is impossible to know from our data whether one or both of these TFs are important in the celastrol response (notably, they do have different binding patterns *in vivo* in the untreated condition; Supplemental Figure 11B,C). Nevertheless, our regression framework is useful in providing a list of candidate TFs whose binding preferences correlate with aspects of the celastrol response.

### Increased polymerase pausing is broadly associated with transcriptional repression

Promoter-proximal pausing of RNA polymerase is a rate-limiting and independently regulated step in productive transcription^53,54^. Notably, the peaks of paused RNA polymerase at DE genes doubled in height during our time course (Figure 5A). Accordingly, we found that the “log pause index,” or log_2_ ratio of average read depth at the pause peak to that in the proximal gene body, increased by more than 1 (corresponding to a fold-change of more than 2 in the pause index) in DE genes by 160 minutes (Figure 5B). Together, these observations indicate that most DE genes undergo increased pausing after celastrol treatment, suggesting that pause release may be widely inhibited during the celastrol response.

**Figure 5.**
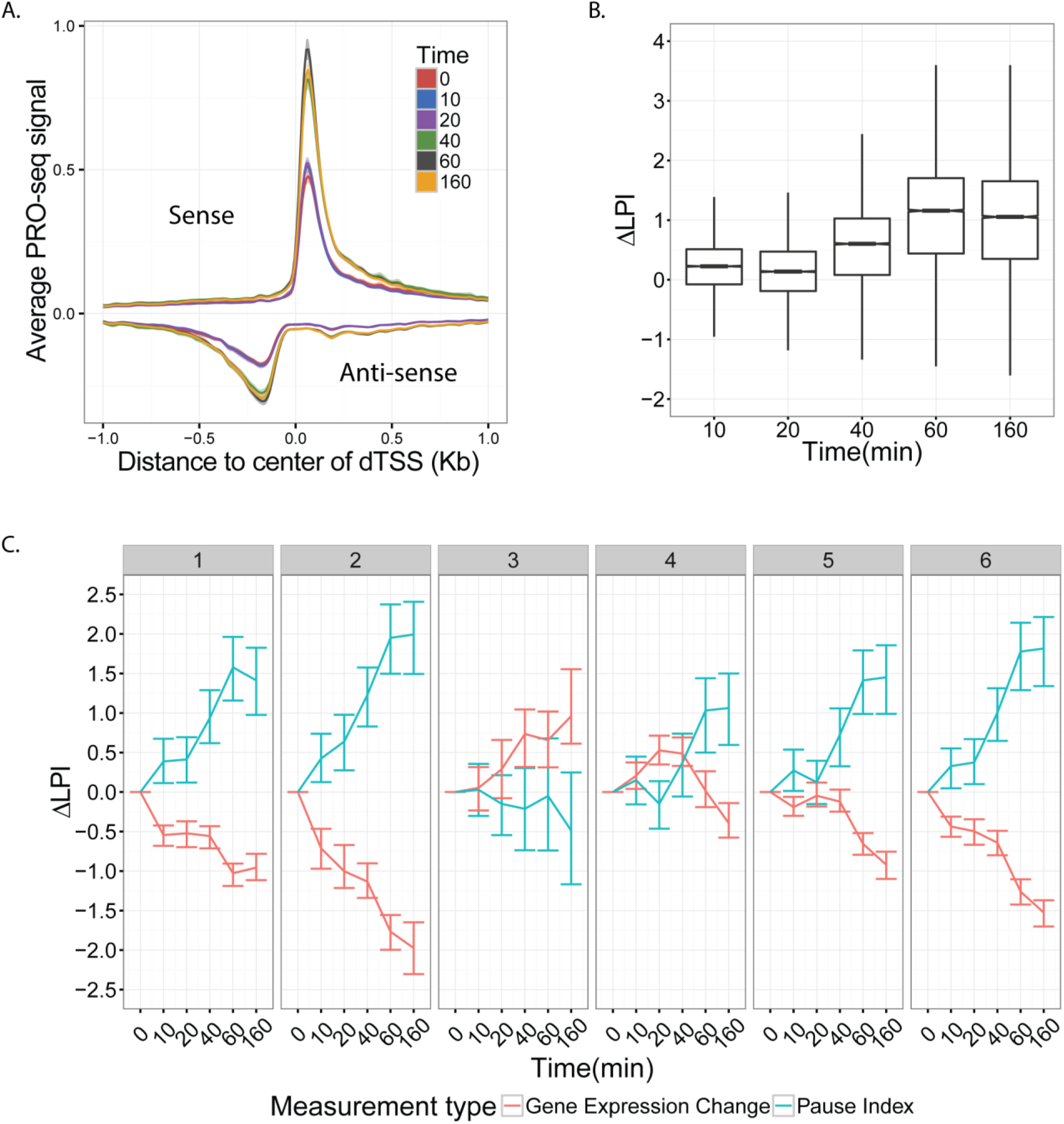
Increased promoter-proximal pausing is associated with transcriptional repression in the response to celastrol. **(A)** Mean PRO-seq signal at promoters for all active genes, grouped by time point and oriented with respect to the direction of transcription of the gene. X-axis represents distance to the center of the divergent transcription start site (see Methods). Intervals around each line represent 95% confidence intervals obtained by bootstrap sampling. Notice the general increase in the height of the pause peaks with time. **(B)** The distribution of changes in the log fold index with respect to the untreated condition (ΔLPI; see Methods) for all active genes at each time point. **(C)** The distribution of ΔLPI for all DE genes (FDR ≤ 0.01) by cluster and time-point. Notice that all clusters show an increase in the pause index with time, except for cluster three.

To see if particular expression patterns were associated with changes in pausing, we separately examined the changes in log pause index during the time course for each of our six gene expression clusters. Interestingly, we found that pausing increased in all clusters with the exception of cluster #3 (Figure 5C), the only strongly up-regulated cluster (see Figure 3A), where it remained essentially unchanged. Thus, changes in the log pause index are generally negatively correlated with changes in expression across clusters. This observation suggests that decreases in the release of paused Pol II to productive elongation could contribute to increased pausing and, hence, to down-regulation of transcription, while the absence of such an effect (in cluster #3) might permit up-regulation of transcription^15,56^. As cluster #3 is strongly associated with the HSF1 response, this finding is consistent with previous reports that HSF1 regulates transcription by increasing the rate of release of paused RNA polymerase into productive elongation^15^.

### Enhancers show similar functional associations and pausing patterns to genes

Previous studies have shown that putative enhancers are divergently transcribed, producing nascent RNAs that can be detected via PRO-seq^17,18,57^. Using the dREG prediction method^58^, we identified 25,891 apparent divergent transcription start sites (dTSS) from our PRO-seq data, pooling data across time points. Based on the distance from nearest annotated TSS, we classified 7,334 of these dTSS as likely transcribed enhancers, 15,941 as likely promoters, and the remaining 2,616 as ambiguous. For validation, we examined ChIP-seq data from ENCODE for untreated K562 cells, and found, as expected, that enhancer and promoter classes were both strongly enriched for acetylation of histone H3 at lysine 27 (H3K27ac), and that the promoter class was more strongly enriched for RNA polymerase and trimethylation of histone H3 at lysine 4 (H3K4me3) (Figure 6A). The enhancer class also showed moderate enrichment for monomethylation of histone H3 at lysine 4 (H3K4me1). These observations confirm that PRO-seq serves as an efficient single-assay approach for characterizing both transcribed enhancers and protein-coding genes^58^.

**Figure 6.**
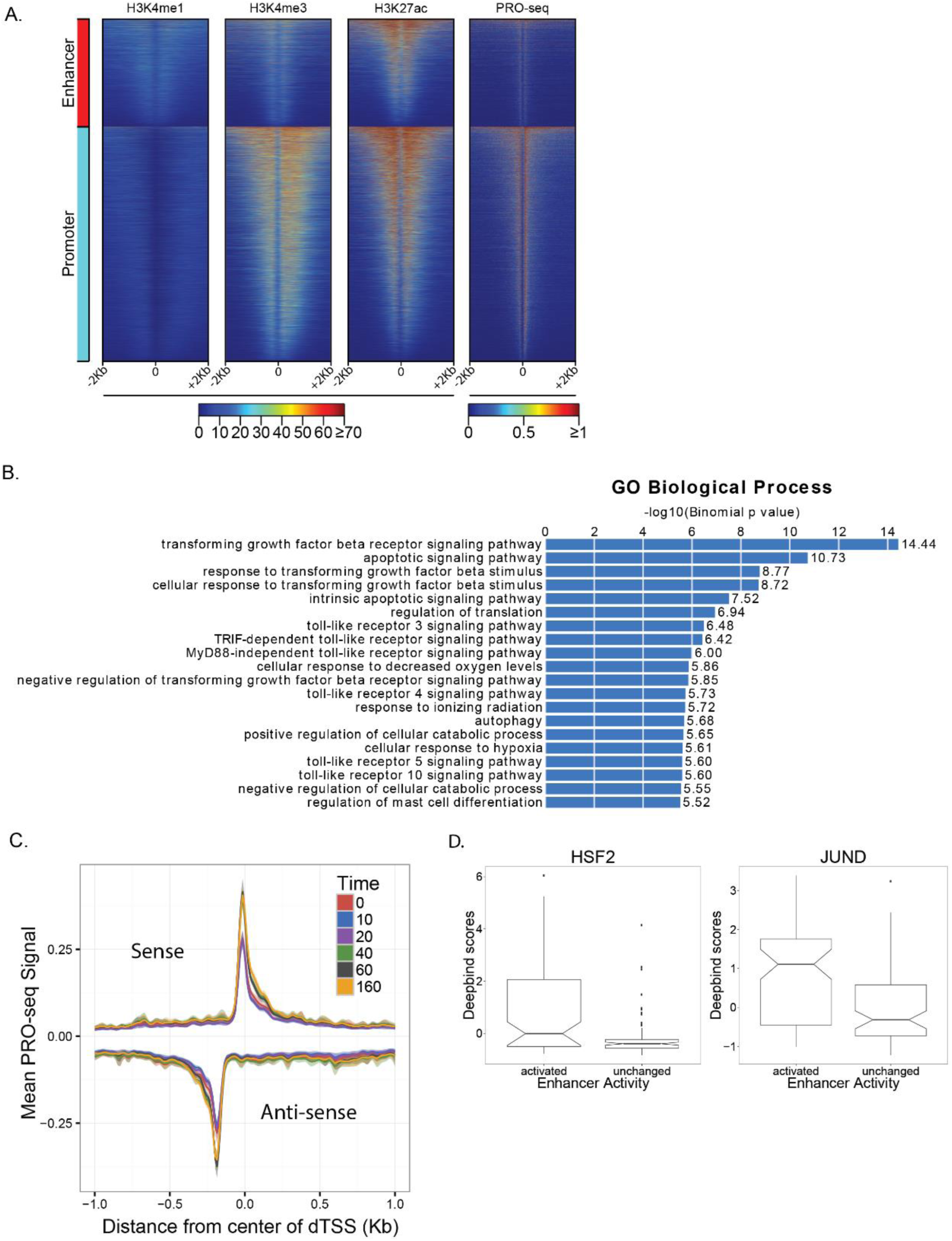
Response to celastrol at predicted transcribed enhancers. **(A)** Divergent transcription start sites (dTSS) that were classified by distance-based rules as likely enhancers or promoters show distinct patterns of histone marks in untreated K562 cells. Shown are H4K4me1 (enriched at enhancers), H3K4me3 (enriched at promoters), and H3K27ac (enriched at both). PRO-seq read counts are shown for comparison. X-axis is oriented by direction of transcription of nearest gene. **(B)** Gene Ontology (GO) biological processes associated with DE transcribed enhancers using GREAT^59^. Bar plot represents −log10 p-values for enrichment, with numerical fold enrichments indicated at right. **(C)** Metaplot of PRO-seq signal at all enhancers, centered on the dTSS, per timepoint.Units of PRO-seq signal are average numbers of reads per 10bp bin. Intervals around each line represent a 95% confidence interval obtained by bootstrap sampling.

To better understand the role of the non-coding regulatory elements in the celastrol response, we further examined 1,479 (~20%) of the 7,334 dTSS-based enhancers that were classified as differentially transcribed. We attempted to find functional enrichments for potential target genes of these differentially transcribed enhancers using the Genome Regions Enrichment of Annotations Tool (Ver. 3.0) (GREAT^59^; see Methods). GREAT identified enrichments for processes relating to apoptosis, translational regulation, and responses to various environmental stresses (Figure 6B), in general agreement with our analysis of DE genes. We also found that our set of putative enhancers displayed an accumulation of paused polymerase after celastrol treatment (Figure 6C)(Supplemental Figure 12). Although the functional significance of pausing at enhancers is unknown, this observation suggests that global shifts in pause levels at genic TSS are also reflected at enhancers.

Lastly, we sought to determine which TFs influenced activity at a relatively small group of 480 enhancers that showed little activity at 0 minutes but greatly increased activity by 160 minutes. We compared DeepBind scores for these activated enhancers with those for non-DE enhancers that had similar absolute expression levels and found six TFs with significantly elevated scores in the activated enhancers: HSF2, JUND, FOSL2, MAFK, STAT3, and THRA (Supplemental Figure 13; Figure 6D). Of these TFs, HSF2, was also associated with increased expression in genes, while JUND and FOSL2 are subunits of AP-1, a TF previously found to regulate cellular growth and senescence^60^. Because these TFs were identified simply based on their sequence preferences, TFs with similar motifs are also potential regulators. For example, it is possible that JUN, whose expression increases over the time course and which is known to be activated by HSF1^61^, is actually responsible for the apparent association with JUND, which does not appear to be activated. Together, these results demonstrate that PRO-seq can be used to detect transiently activated enhancers and identify TFs that may drive the enhancer response.

## Discussion

This study represents the first genome-wide assessment of the immediate transcriptional effects of celastrol, including transcribed regulatory elements as well as genes, shedding light on some of the possible primary targets and mechanisms of action of this potent therapeutic compound. We find that celastrol treatment results in pervasive transcriptional down-regulation, with nearly half of expressed genes being down-regulated within 160 minutes. A much smaller group of genes, roughly 10% of those expressed, are up-regulated during the same time interval. By analyzing the sequences nearby transcription units, we were able to identify several transcription factors whose binding patterns partially explain these transcriptional responses. We also observed a clear impact from celastrol on polymerase pausing at both genes and enhancers, which is negatively correlated with changes in transcriptional activity. While there are limits to what can be learned from PRO-seq data alone, we have shown that when these data are collected at relatively high temporal resolution and analyzed together with other data for the untreated condition, they can provide valuable insights into a multifaceted, multistage cellular response to a transcriptional stimulus.

We find that celastrol treatment generally induces a similar response to heat shock, consistent with previous reports^19,30^. Heat-shock has also been observed to induce widespread down-regulation in mammalian cells^15,37^. Moreover, many of the same genes that are up-regulated upon celastrol treatment are also bound by HSF1 after heat shock or participate in heat-shock pathways. Finally, sequences associated with HSF1 binding are associated with increased gene expression, according to our regression analysis. Together, these findings suggest that HSF1 is activated soon after celastrol treatment, whereupon it activates a large group of genes.

These observations suggest that, in part, the transcriptional response to celastrol may simply be a general cellular stress response. Cellular stress responses are known to affect a broad range of cellular functions, including cell cycle arrest, transcription of molecular chaperones, activation of DNA damage repair pathways. removal of irretrievably damaged macromolecules, and apoptosis upon severe damage^62^, and they are relevant in many diseases, including cancer^63^, proteotoxic diseases^31^, and autoimmune disorders^64^. Previous studies have investigated cellular stress responses at the delayed transcriptional^15,65^, post-transcriptional^66^, translational^40^, and post-translational levels^67,68^. Together with similar studies of heat shock^15^, our study is valuable in illuminating features of the early transcriptional response to stress.

Nevertheless, we observed differences between the heat shock and celastrol responses. For instance, we found that binding sites for SRF, a transcription factor associated with HSF1/2- independent up-regulation after heat-shock, are enriched in the core promoters of genes having elevated expression after 40 minutes^15^. By contrast, previous findings for heat shock^15^ have indicated that the positive effect on gene expression occurs as early as 2.5 minutes after treatment, suggesting that there may be some kinetic differences between the celastrol and heat-shock responses. Our analysis also suggests that the loss of binding by MYC-MAX may be responsible, in part, for the broad transcriptional repression within 20 minutes of celastrol treatment. This finding is supported by previous studies showing that celastrol directly inhibits MYC-MAX functionality^49^. Since MYC-MAX is a strong transcriptional activator and is bound at over 6,000 promoters of active genes in K562, its inhibition may be an important contributing factor to widespread down-regulation after celastrol treatment^69,70^. To our knowledge, MYC-MAX inhibition has not been reported to be important in the heat-shock response, but if it is, this inhibition is likely to be due to a different mechanism.

A major strength of our experimental approach is that it allows us to observe transient as well as sustained transcriptional responses. For example, we found that E2F4 was quickly down-regulated after celastrol treatment, reducing in expression by half within 20 minutes, but, nevertheless, target genes of this transcription factor were increasingly repressed later in our time course (between 60 and 160 minutes). These observations suggest that the apparent increased activity of E2F4 during the celastrol response may have a non-transcriptional basis; for example, perhaps pre-existing E2F4 is post-translationally modified to become more active. Interestingly, previous studies have shown that celastrol inhibits CDK4, and that CDK4 overexpression disrupts E2F4 DNA-binding ability^71,72^. Thus, it is possible that celastrol treatment frees up E2F4 to bind DNA, which in turn could contribute to cell-cycle arrest^47,48^.

Another advantage of our densely sampled PRO-seq time course is that it allows us to measure changes in promoter-proximal RNA polymerase pausing. We observed that pause indices increased by more than two-fold at differentially expressed genes during our time course. We also found that increase pausing was associated with decreased transcription in genes, as previously reported for heat-shock conditions^15^, although we did not observe a converse association between decreased pausing and up-regulation of genes. A possible mechanism behind this increased pausing is the disruption of the MYC-MAX complex, which has been shown to recruit P-TEFb, which in turn facilitates pause release^73^. This mechanism could in principle affect down-regulated genes only, for example, if up-regulated genes recruit P-TEFb independently of MYC-MAX (e.g., through binding of HSF1^74^).

While it is possible that increased pausing causes decreased expression, by limiting productive elongation and therefore reducing transcription levels, an alternative possibility is that decreased transcriptional activity across many genes results in increased availability of free Pol II, some of which ends up being loaded on promoters and coming to rest at pause sites^15^. In other words, the negative correlation between pausing and expression could be explained by causality in either direction, or perhaps in both directions. Additional experiments will be needed to establish the causal basis of these correlations. In any case, our observations suggest that changes in pausing are widespread and broadly associated with transcriptional repression, and therefore may play an important role in the celastrol response.

## Materials and Methods

### Celastrol treatment

K562 cells were cultured at 37 °C in RPMI media (Gibco) containing 10% FBS (Gibco), Pen Strep (Gibco) and 2 mM L-Glutamine (Gibco). Biological replicate cell cultures were prepared as follows: after thawing K562 cells and seeding a fresh culture, cells were split into two separate flasks, which would remain separated through six passages and expansions until treatment and collection for preparation of PRO-seq libraries. Cells from each expanded replicate were seeded onto six 30-mL dishes (one for each time point) at a density of 5x10^5^ cells/mL and then incubated for an additional doubling cycle (~20 hrs). For treatments, fresh celastrol was dissolved in DMSO at a final concentration of 20 mM. Celastrol-treated samples received celastrol (Sigma) at a final concentration of 3 μΜ, whereas untreated (0-minute) samples received an equivalent volume of DMSO. Cells remained in culture dishes in the incubator during the time course. Time-course treatments were carried out in reverse order so that all samples would be collected at the same time (starting with 160-minute time point and ending with the untreated).

### Cell permeablization and PRO-seq

Samples were then prepared for precision run-on reactions by subjecting cells to permeablizing conditions. Briefly, cultures were spun down and resuspended in ice cold 1xPBS. Samples were spun again and washed in 5 mL wash buffer (10 mM Tris-Cl, pH 7.5; 10 mM KCl; 150 mM sucrose; 5 mM MgCl_2_; 0.5 mM CaCl_2_; 0.5 mM DTT; 1x Protease inhibitor cocktail (Roche); 20 units RNase inhibitor (SUPERase In, Invitrogen)). Cell pellets were then resuspended in permeablization buffer (10 mM Tris-Cl, pH 7.5; 10 mM KCl; 250 mM sucrose; 5 mM MgCl_2_; 1 EGTA; 0.05% Tween-20; 0.1% NP40; 0.5 mM DTT; 1x Protease inhibitor cocktail (Roche); 20 units RNase inhibitor (SUPERase In, Invitrogen)) and left on ice for 5 minutes. Cells were checked for penetration by trypan blue to assess permeability (~99% permeable). Cells were then washed two times in 5 mL wash buffer before being resuspended in 200 μL storage buffer (50mM Tris-Cl, pH 8.3; 40% glycerol; 5 mM MgCl_2_; 0.1 mM EDTA; 0.5 mM DTT). A one-to-fifty dilution was prepared using 2 μL of each sample and used to take OD_600_ measurements. All samples were then diluted to an equal density (OD_600_ = 0.181) in a final volume of 110 μL of storage buffer. 5x10^4^ pre-permeabilized S2 cells were then spiked in to each cell count-normalized sample before flash-freezing the permeablized cells and storing them at −80 °C.

Stored permeable cells with spike-ins were thawed on ice and each sample was subjected to the precision run-on protocol^75^. Run-on reactions incorporated only biotinylated NTPs with no un-modified NTPs. All libraries were subjected to nine cycles of PCR amplification before size selection and gel purification.

### Read mapping

All filtered reads were removed from each fastq file, then cutadapt (v1.9.1) was run with the following options:

> cutadapt -a TGGAATTCTCGGGTGCCAAGG -m 15

to remove the Illumina adapters and discard all remaining reads that were less than 15bp in length. All reads were then trimmed to 34bp in length using fastx_timmer (v0.0.13.2) to avoid biasing read mapping away from gene promoters. The trimmed reads were then aligned to the joint hg19/bdgp6 genome using the STAR aligner (v2.4.0i). Reads aligning to hg19 and bdgp6 were then separated and bigwigs were created by converting each read to a single count at its 5’ end.

### Detection and resolution of dTSS

dREG was run on each sample as described previously^58^, producing a set of genomic intervals corresponding to predicted divergent transcription starts sites (dTSS). These initial dREG calls had fairly coarse resolution, ranging in size from several hundred to thousands of bases. We therefore applied a heuristic scanning method to identify one or more higher-resolution dTSSs within each dREG call. Briefly, this method involved sliding a window along a dREG interval and considering the relative read counts among three subintervals: a peak, a flank, and center. To identify pairs of divergent peaks, the test was applied simultaneously to each strand in a strand-specific manner, and the results were combined. Specifically, for a scan initiated at base *i*, the center was defined as the interval [*i*, *i* + 110), the shoulder as [*i* − 50, *i*), and the flank as [*i* − 250, *i* − 150) (Supplemental Figure 2). Three one-sided binomial tests were performed, testing that there are fewer reads in the center than the flank, the center than the shoulder, and the flank than the shoulder. The sum of the resulting six negative log p-values (three for each strand) then became the per-base score. The best scoring window in a dREG region was taken as a dTSS. In addition, up to two other dTSSs were called if their score exceeded 20.

### Classifications of dTSS

dTSS were classified as either enhancers or promoters based on their relative distance from the set of all TSSs annotated in GENCODE v19. To classify each dTSS, the following rules were applied: (1) if the dTSS was greater than 1 kb, and at most 1 Mb, away from the nearest annotated promoter, it was classified as an enhancer; (2) if the dTSS was within 200 bp of an annotated promoter, or it was within 1 kb of an annotated promoter and it was the closest dTSS to the promoter, it was classified as a promoter; (3) if the dTSS was between 600bp and 1 kb away from the nearest annotated promoter, and not the closest dTSS to the promoter, It was classified as an enhancer; (4) otherwise, the dTSS was classified as unknown.

### Selection of active transcripts in K562 cells

Selection of transcripts was performed by a new program, called TuSelector. First, a list of potential transcripts was obtained from GENCODE v19. The genic regions and data were partitioned into 100bp intervals. For each gene, a set of coarse-grained overlapping transcript models was created, where for each transcript model and interval, the interval was assigned to the transcript model if, and only if, it overlapped the corresponding annotated transcribed region by more than 50% at the nucleotide level. Next, the PRO-seq read counts in each 100 bp interval were summarized by a 1 if there were reads aligned to the interval or a 0 otherwise. TuSelector computed a likelihood for each of the possible coarse-grained transcript models at a given gene, as follows:

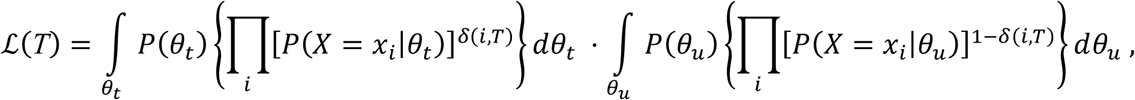

where *T* is the transcript model, the products range across genomic intervals *i*, *x_i_* is the summary of the data in interval *i* (0 or 1), *δ*(*i*, *T*) is an indicator function that takes value 1 when interval *i* is included in *T* and 0 otherwise, *X* is a Bernoulli random variable, and *θ_t_* and *θ_u_* are the parameters for this random variable in the transcribed and untranscribed states, respectively. *P*(*θ_t_*) is assumed to be uniform over the interval (0.3, 1), and *P*(*θ_u_*) is assumed to be uniform over the interval (0.01, 0.03). In practice, we discretized *θ_u_* into segments of size 0.01 and *θ_t_* into segments of size 0.05, and approximated the integrals with finite sums. Finally, in addition to the annotated transcripts, we considered a competing model representing a completely untranscribed gene.

TuSelector was run separately for each replicate and time point, and potentially produced discordant transcript calls across these runs. Therefore, we selected at most one “consensus” transcript model per gene for use in further analysis, as follows. To be considered a consensus call, TuSelector had to identify the same transcript model at least 80% of the time with at least 50% of replicate pairs both having the same transcript call. Two transcript models were considered “the same” if their endpoints differed by less than 500bp. If no transcript model met these criteria, the gene was not considered in further analysis.

### Estimating expression and detecting differentially expressed genes

For all active, protein-coding transcripts, reads were taken from up to the first 16 kb of the gene, minus the first 500bp to avoid an influence from promoter-proximal pausing. This strategy allowed us to focus on the most recent transcription at each time point and avoid averaging over time. The maximum interval of 16 kb was based on a minimum interval between time points of 10 minutes and an average polymerase transcription rate of ~2 kb/min, minus a few kilobases of “padding”. Any genes that were shorter than 700 bp were removed from the analysis. A size factor for each sample was obtained by taking the number of spike in reads per sample divided by the median number of spike in reads per sample. To estimate expression of transcriptional enhancers, reads were taken from 310 bases (assuming a 110 base spacing between dTSS as reported by Core et al. plus 100bp to either side) centered on the dTSS. Both sets of read counts were fed jointly into DESeq2, and enhancers and genes were subsequently separated for further analysis. An enhancer or gene was called as DE with an FDR ≤ 0.01 using a likelihood ratio test.

### Clustering differentially expressed genes

Gene expression log-fold changes were computed relative to the untreated (zero-minute) time-point using the DESeq-based estimates of absolute expression (rlog values). All DE genes were then fed into the autoregressive clustering program EMMIX-WIRE using default settings. Likelihood values for between two and ten clusters were computed. We selected six clusters as a value at which the increases in likelihood with the number of clusters began to decline. To check for the robustness of our selection we repeated our analyses with five and seven clusters and found that they were not highly sensitive to the cluster number.

### Computing functional enrichment for gene clusters

Reactome (v52) was used to assign genes to functional categories. Genes that were not annotated in Reactome were removed. The background set for all enrichments was the set of DE genes present in Reactome. Odds ratios were computed per cluster (*c*) and pathway (*p*) as:

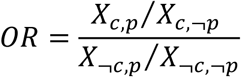

where *X* represents a count and *c* and ¬*c* denote the sets of genes in, and not in, cluster *c,* respectively, and, similarly, *p* and ¬*p* denote the sets of genes in, and not in, pathway *p*. An empirical null distribution of odds ratios was computed by randomly shuffling the gene assignments to pathways 100,000 times. *P*-values were then computed from this distribution and the Benjamini-Hochberg procedure was applied to estimate false discovery rates (FDRs).

### Characterizing genic regulation

ChIP-seq data was downloaded from the ENCODE website (https://www.encodeproject.org) in narrowPeak format (optimal idr) on Sep. 30^th^, 2016. Scores for each gene-TF pair were computed by taking the peaks with the maximum signal that intersected [-200,+500] around the promoter. Deepbind v0.11 was run over [-200,+500] around the promoter with standard settings using all non-deprecated motifs for DNA binding proteins. To analyze TFs that may be involved in different regulatory patterns we linearly regressed genic expression (as estimated by DESeq2) against scores from ChIP-seq or DeepBind with time point specific coefficients for each TF and a time agnostic, gene-specific coefficient to capture the fixed effect of unmodeled regulation^46,76^. In this framework, the expected expression of a given gene *i* at time *j* is expressed as

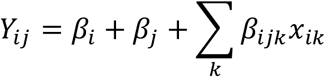

where *β_i_* is the gene-specific expression bias term, *β_j_* is a time-specific bias term, and *β_ijk_* is the coefficient for the time-point specific effect of a TF *k.* Standard deviations for each coefficient were estimated via 1000 bootstraps. Finally, the list of TFs was filtered to keep those with FDR ≥ 0.01 in at least one time point and with a maximal change between any two coefficients in the 90^th^ percentile. This procedure selected for TFs having an effect that was both statistically significant and of large magnitude.

### Identification and analysis of genic pause peaks

To locate pause peaks, we scanned each active transcript (see above) greater than 1 kb in length in the region of the annotated TSS ([TSS − 200, TSS + 200]), taking the number of reads in a 50-bp sliding window, with a sliding increment of 5 bp. The window with the largest number of reads in the untreated condition (0-minute time point) was designated as the pause peak. To compute a log2 pause index (LPI), we subtracted the DESeq-estimated log2 read count (the “rlog” value) for the gene body from the equivalent DESeq-estimated log2 read count at the peak. Furthermore, to compute changes in this value over time, we subtracted the LPI for the zero time point from the LPI for each subsequent time point; that is, the change in LPI at time *t,* denoted ΔLPI_*t*_, was given by ΔLPI_*t*_ = LPI_*t*_ – LPI_0_. Notice that normalizing changes in the pause peak by changes in the gene body in this way only increases if the number of reads in the peak increases by more than the number of reads in the gene body.

### Estimating expression in enhancers

To prevent contamination from genic transcription, all dTSS previously annotated as enhancers were extended by 1 kb to either side and removed if any part of the extended enhancer was within 5 kb of a gene body. DESeq was used to estimate the transcription level in the enhancer peaks, tails, and the whole enhancer body. The enhancer peak was defined as ±250 bp from the center of the enhancer, the tail was ±400 to ±1000 bp from the center, and the whole enhancer was 0 to ±1000 bp from the center of the enhancer. Read counts were summed from both strands for each region (i.e., peak = plus strand [0,+250]+ minus strand[- 250,0]), and then DESeq2 was used to estimate fold changes. Enhancers were called as strongly activated if they were differentially expressed with FDR ≤ 0.01, rlog(expression at 0 min) ≤ 1, and were in the 90^th^ percentile for fold change between 0 and 160 minutes. To get the same number of similarly expressed non-differentially expressed enhancers, we performed rejection sampling on enhancers that were differentially expressed with FDR > 0.5 using their average expression values between 0 and 160 minutes and probabilities calculated from a kernelized histogram of the activated enhancer’s expression at 160 minutes.

## Acknowledgments

We thank other members of the Siepel, Lis, and Danko laboratories for helpful discussions. This research was supported, in part, by US National Institutes of Health (NIH) grants HG007070 (to J.T.L. and A.S.), GM102192 (to A.S.), and HG009309 (to C.G.D.). The content is solely the responsibility of the authors and does not necessarily represent the official views of the NIH.

## Supplemental Material

**Supplemental Figure 1.**
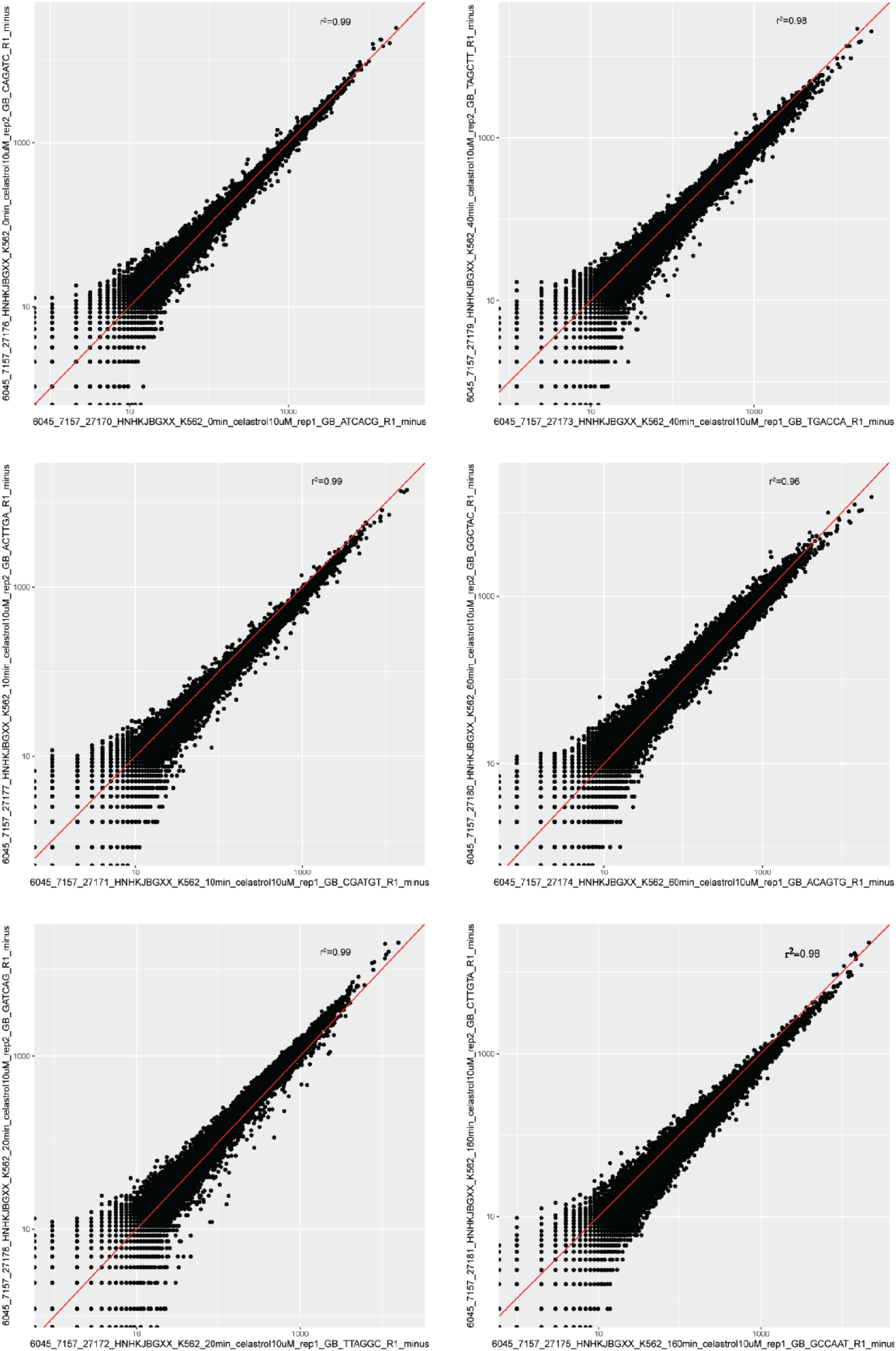
Concordance in read-counts between replicates in gene bodies on the negative strand.

**Supplemental Figure 2.**
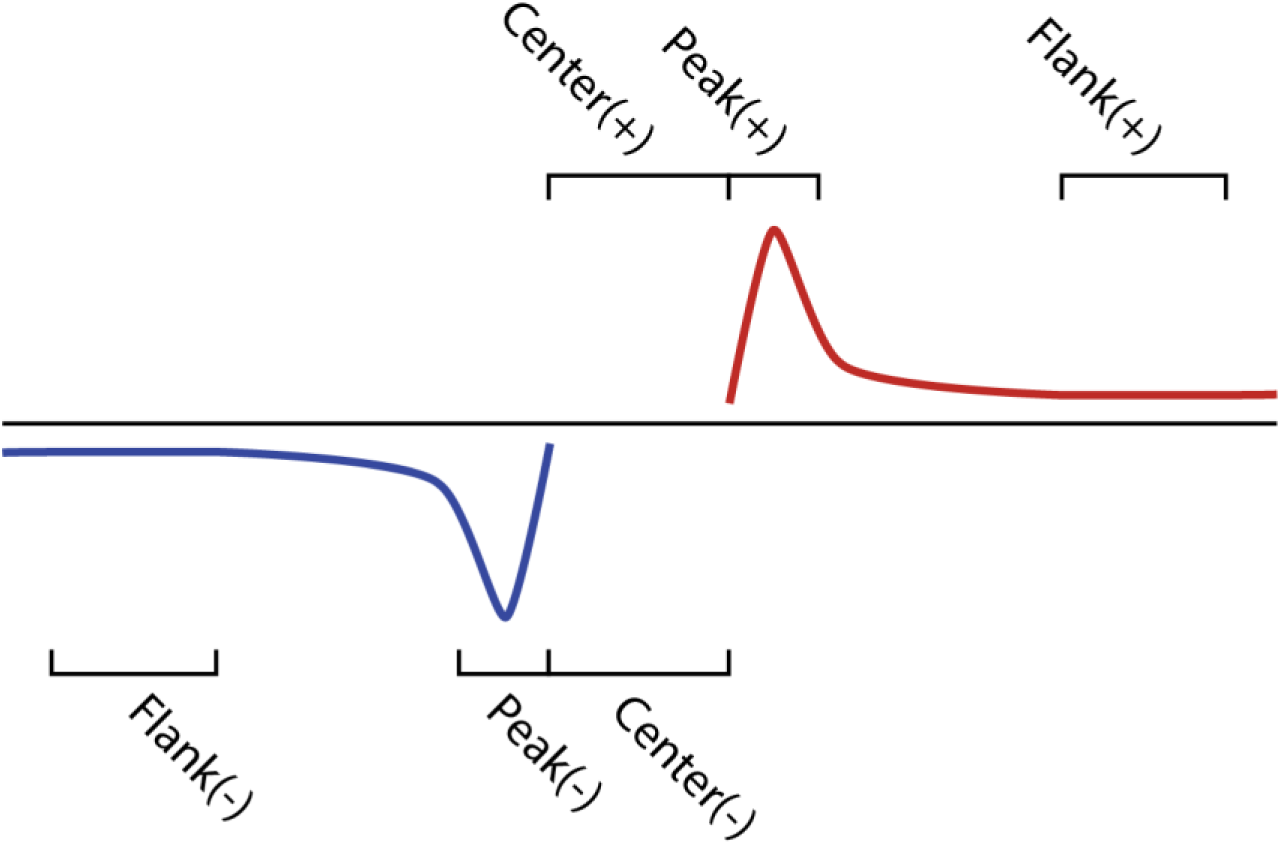
Transcriptional model for refining dREG calls.

**Supplemental Figure 3.**
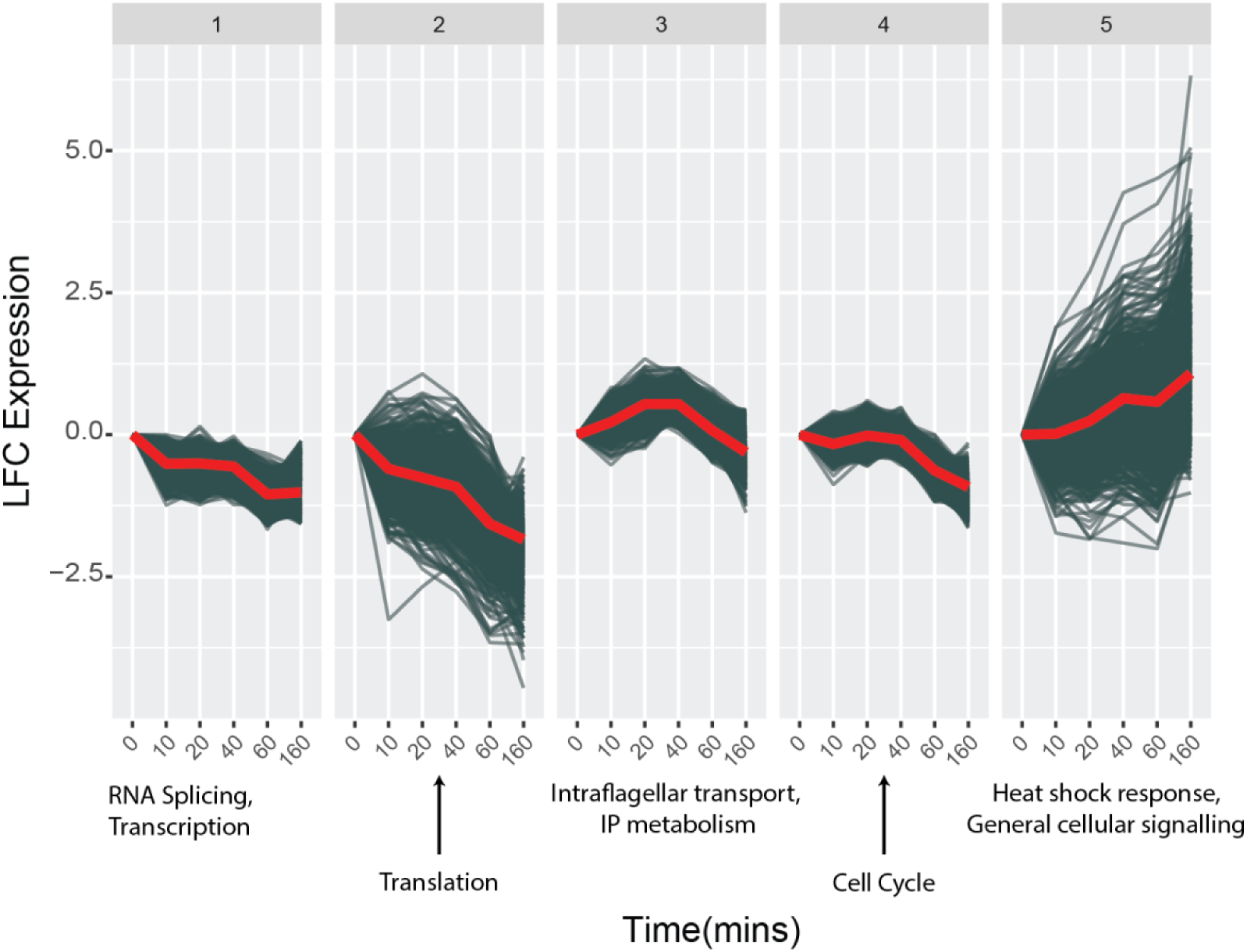
Clustering of DE genes into five clusters and summary of enriched cluster-specific terms.

**Supplemental Figure 4.**
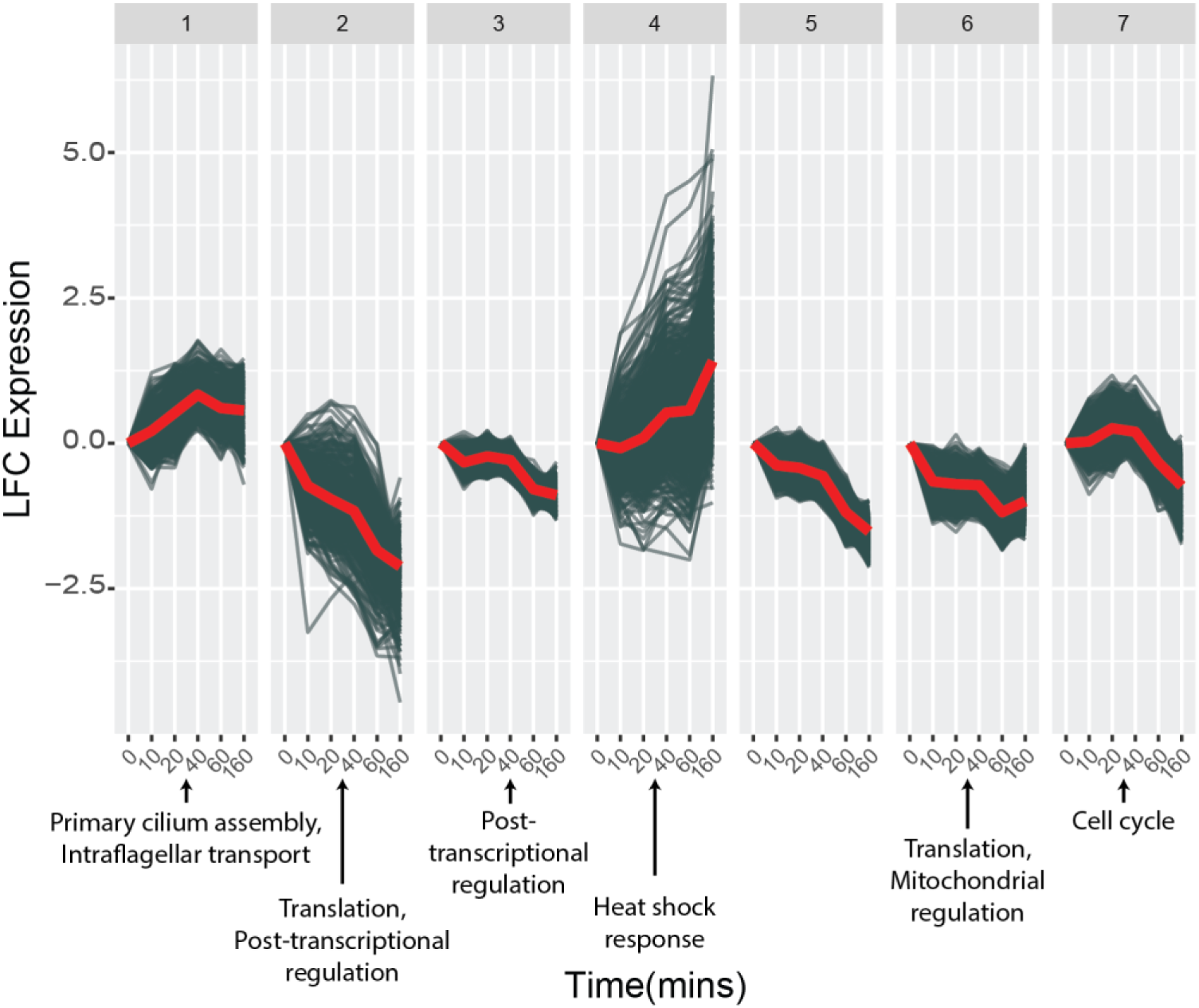
Clustering of DE genes into seven clusters and summary of enriched cluster-specific terms.

**Supplemental Figure 5.**
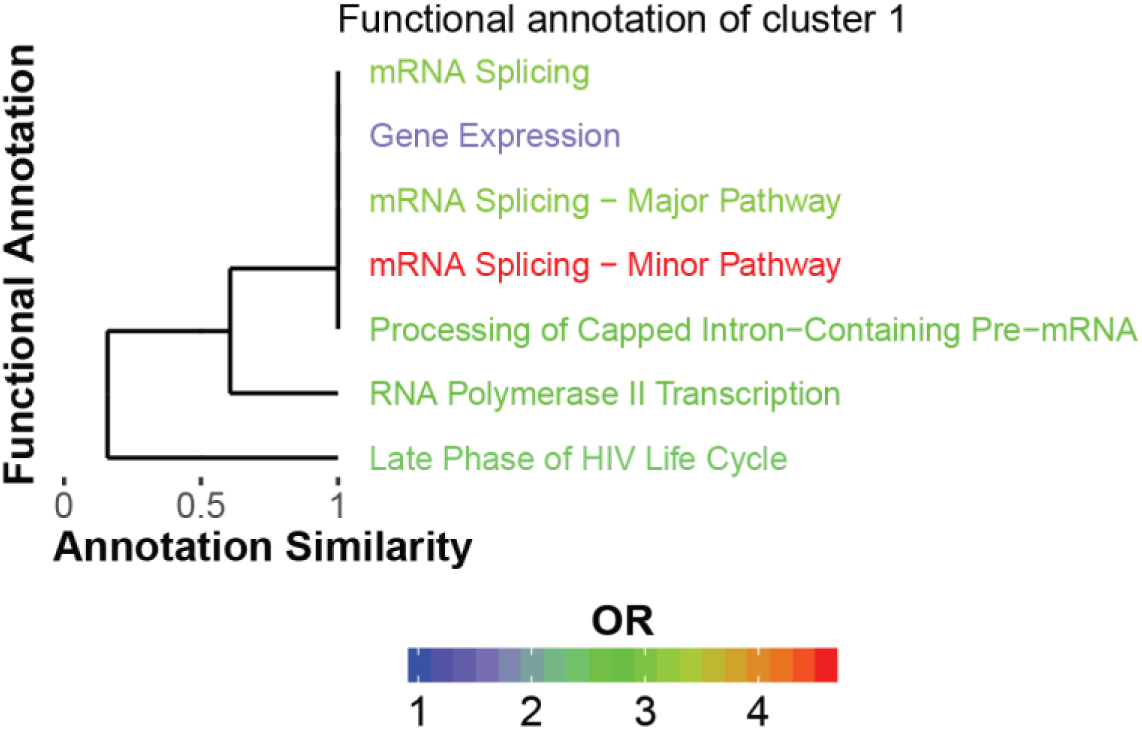
Full set of Reactome terms enriched in Cluster #1 with respect to the other clusters (FDR <= 0.01). Annotation similarity indicates what fraction of genes (based upon the term associated with less genes) are shared between two terms.

**Supplemental Figure 6.**
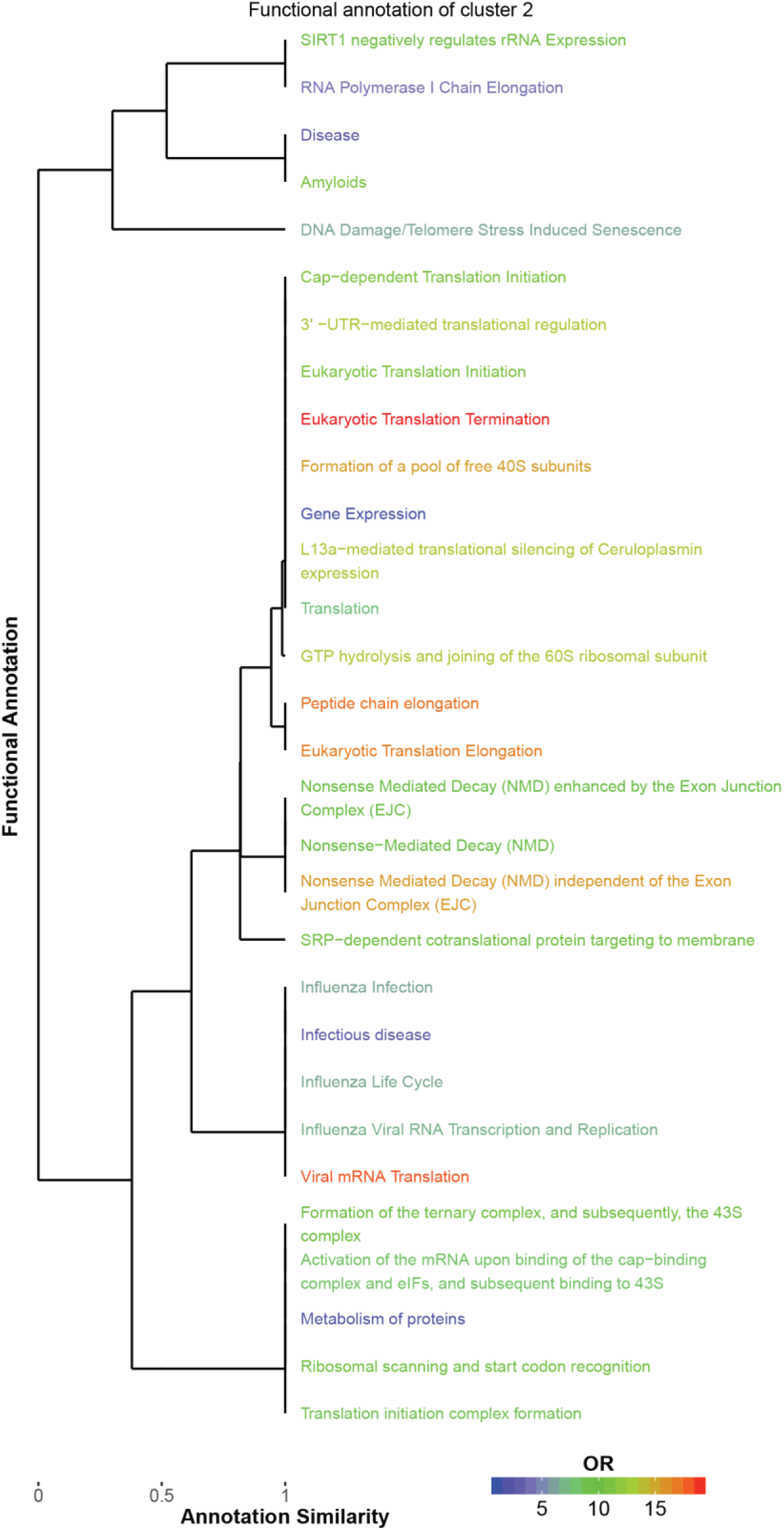
Full set of Reactome terms enriched in Cluster #2 with respect to the other clusters (FDR <= 0.01). Annotation similarity indicates what fraction of genes (based upon the term associated with less genes) are shared between two terms.

**Supplemental Figure 7.**
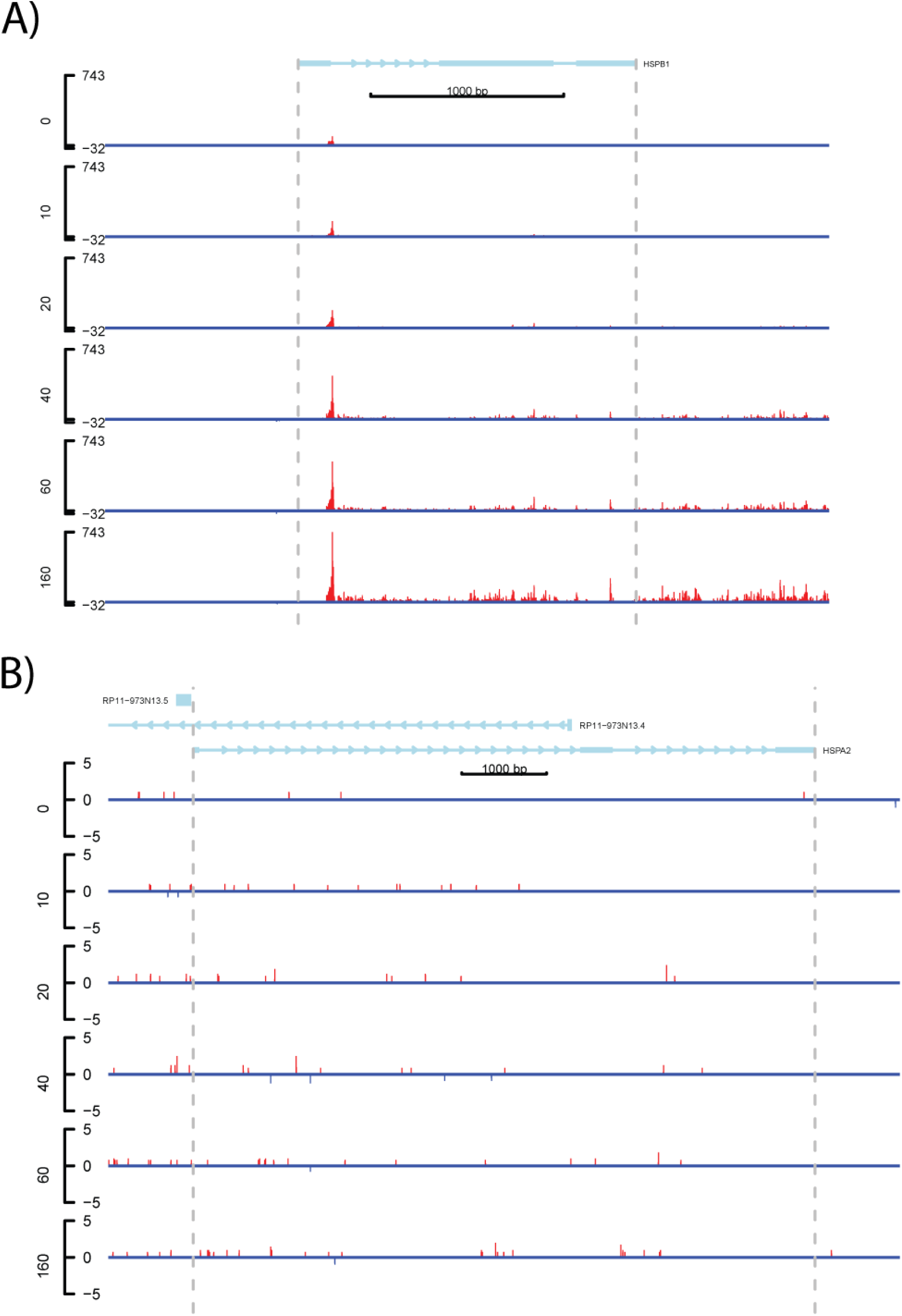
Expression of HSPB1 and HSPA2, key genes in heat-shock induced translational repression. (A) Expression of HSPB1 with each library normalized by size factor and replicates for each time point added together. (B) Expression of HSPA1 with each library normalized by size factor and replicates for each time point added together.

**Supplemental Figure 8.**
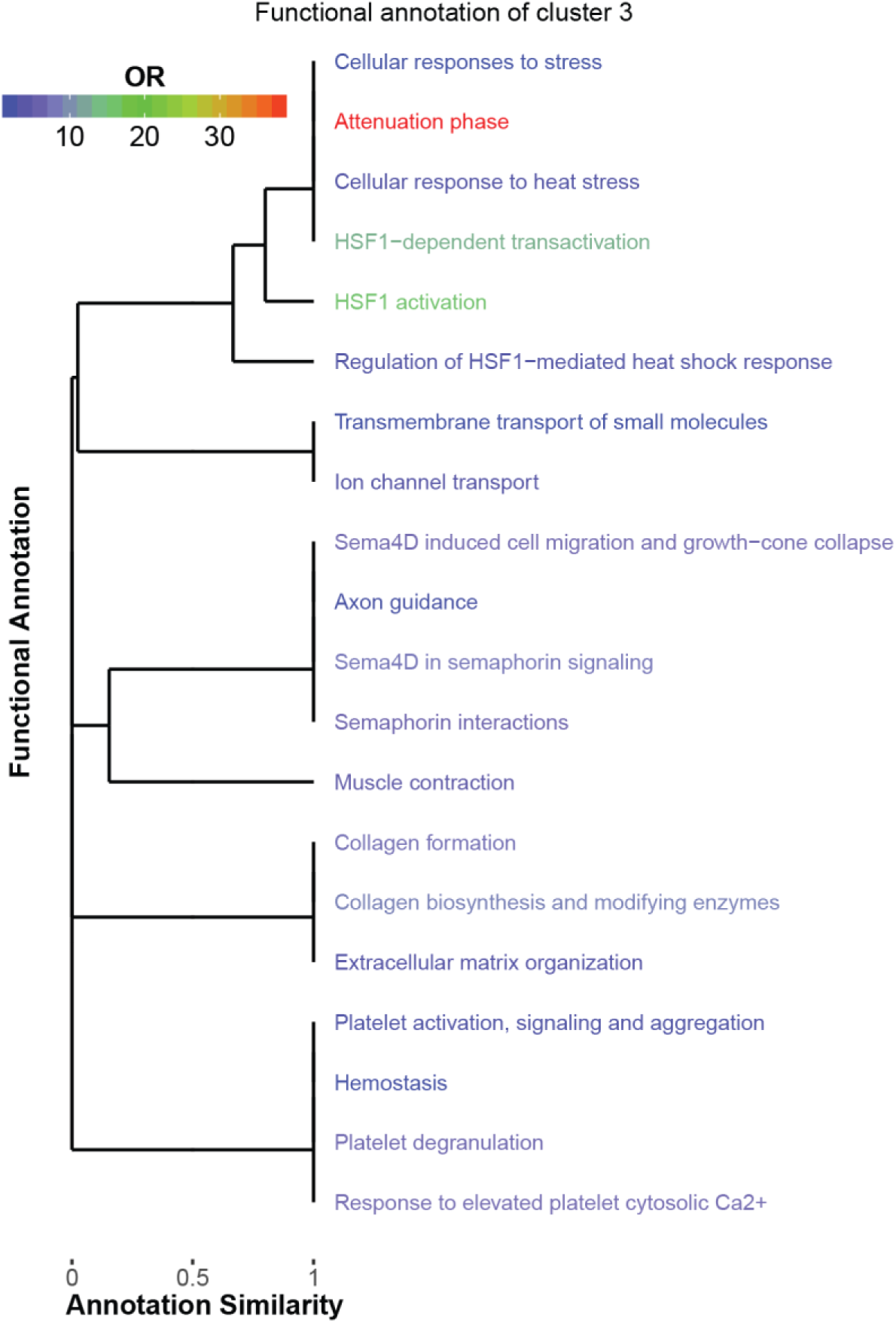
Full set of Reactome terms enriched in Cluster #3 with respect to the other clusters (FDR <= 0.01). Annotation similarity indicates what fraction of genes (based upon the term associated with less genes) are shared between two terms.

**Supplemental Figure 9.**
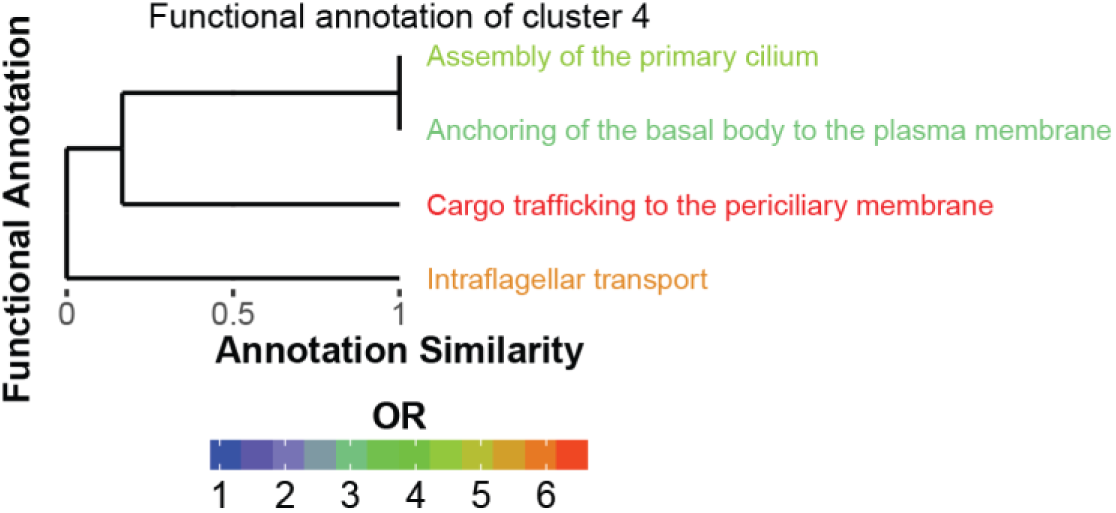
Full set of Reactome terms enriched in Cluster #4 with respect to the other clusters (FDR <= 0.01). Annotation similarity indicates what fraction of genes (based upon the term associated with less genes) are shared between two terms.

**Supplemental Figure 10.**
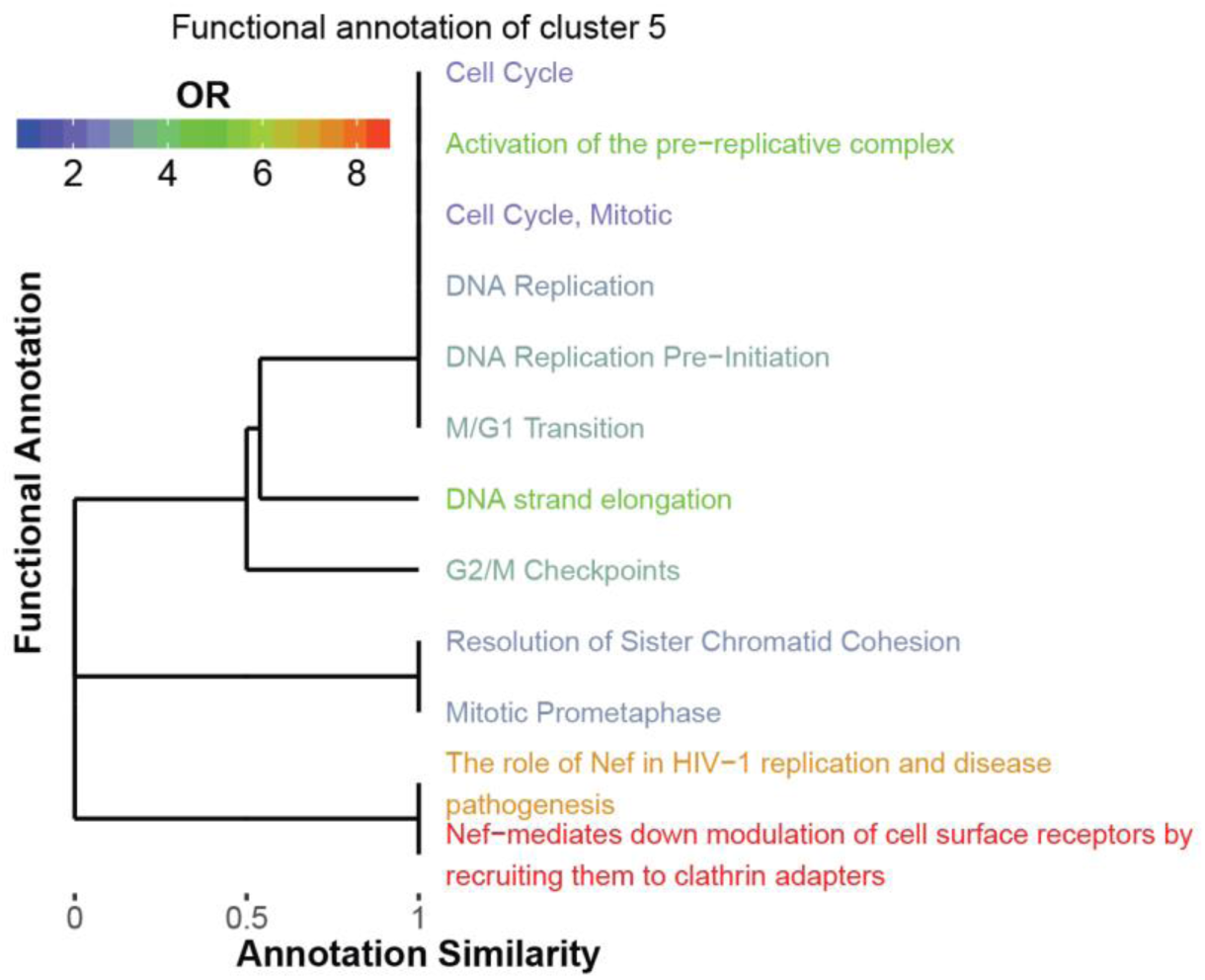
Full set of Reactome terms enriched in Cluster #5 with respect to the other clusters (FDR <= 0.01). Annotation similarity indicates what fraction of genes (based upon the term associated with less genes) are shared between two terms.

**Supplemental Figure 11.**
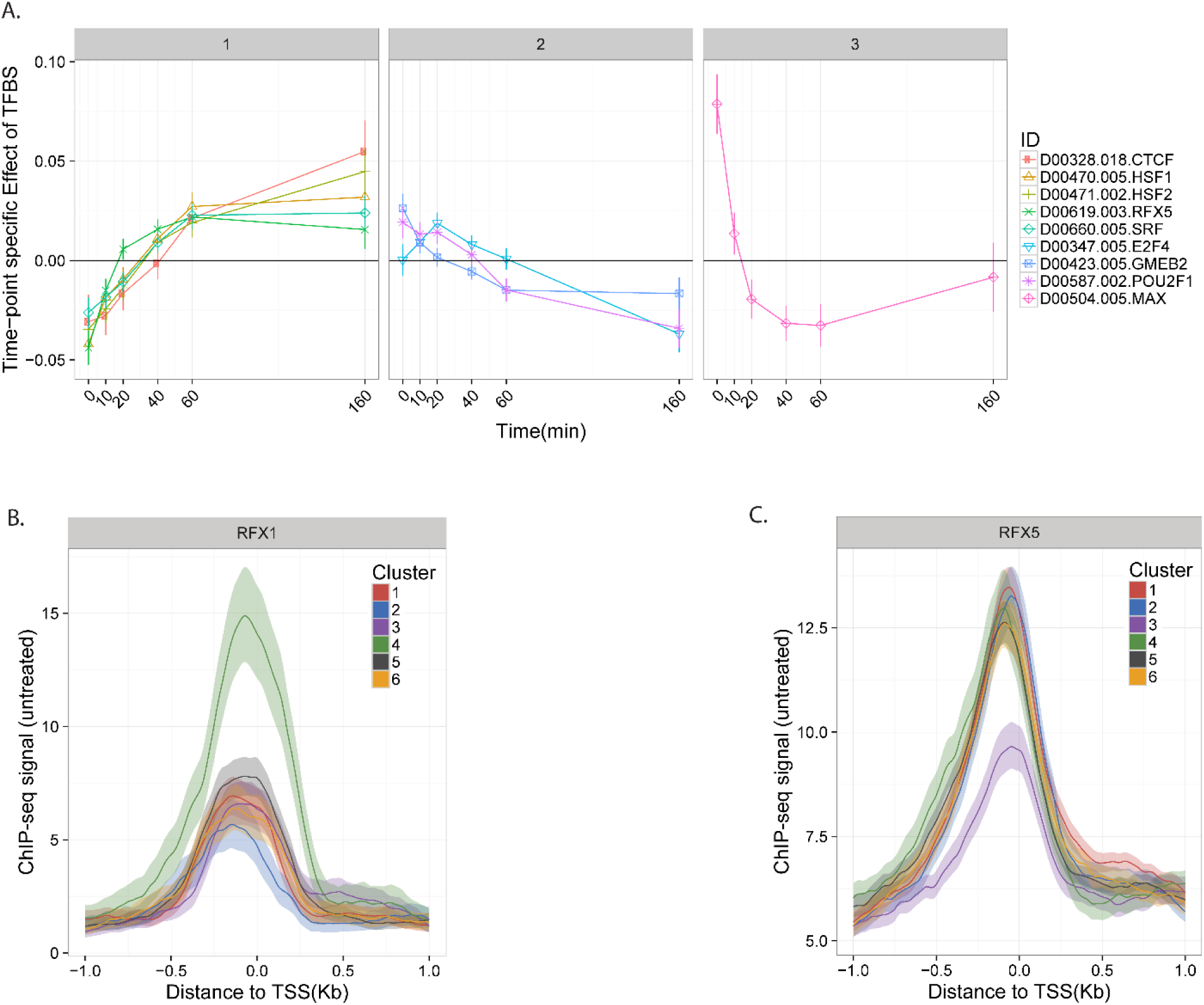
**(A)** Weights from the regression model to predict gene expression from ChIP-seq peak scores in K562. A positive wieght for a TF at given timepoint means that genes with that TF bound showed increased expression relative to those without it. Negative weights mean the opposite. **(B)** Pre-celastrol treatment ChIP-seq signal for RFX1 grouped by clustering based on expression profiles. Each line represents an average over all genes in the cluster in the region of the TSS, with lighter-colored bands representing 95% confidence intervals obtained by bootstrap sampling. **(C)** Same as (B) but for RFX5.

**Supplemental Figure 12.**
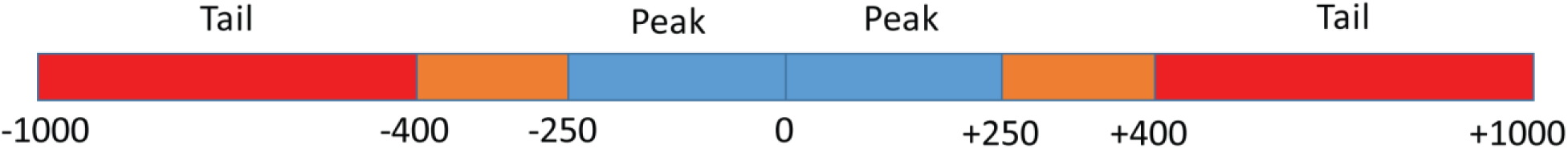
Anatomy of an enhancer.

**Supplemental Figure 13.**
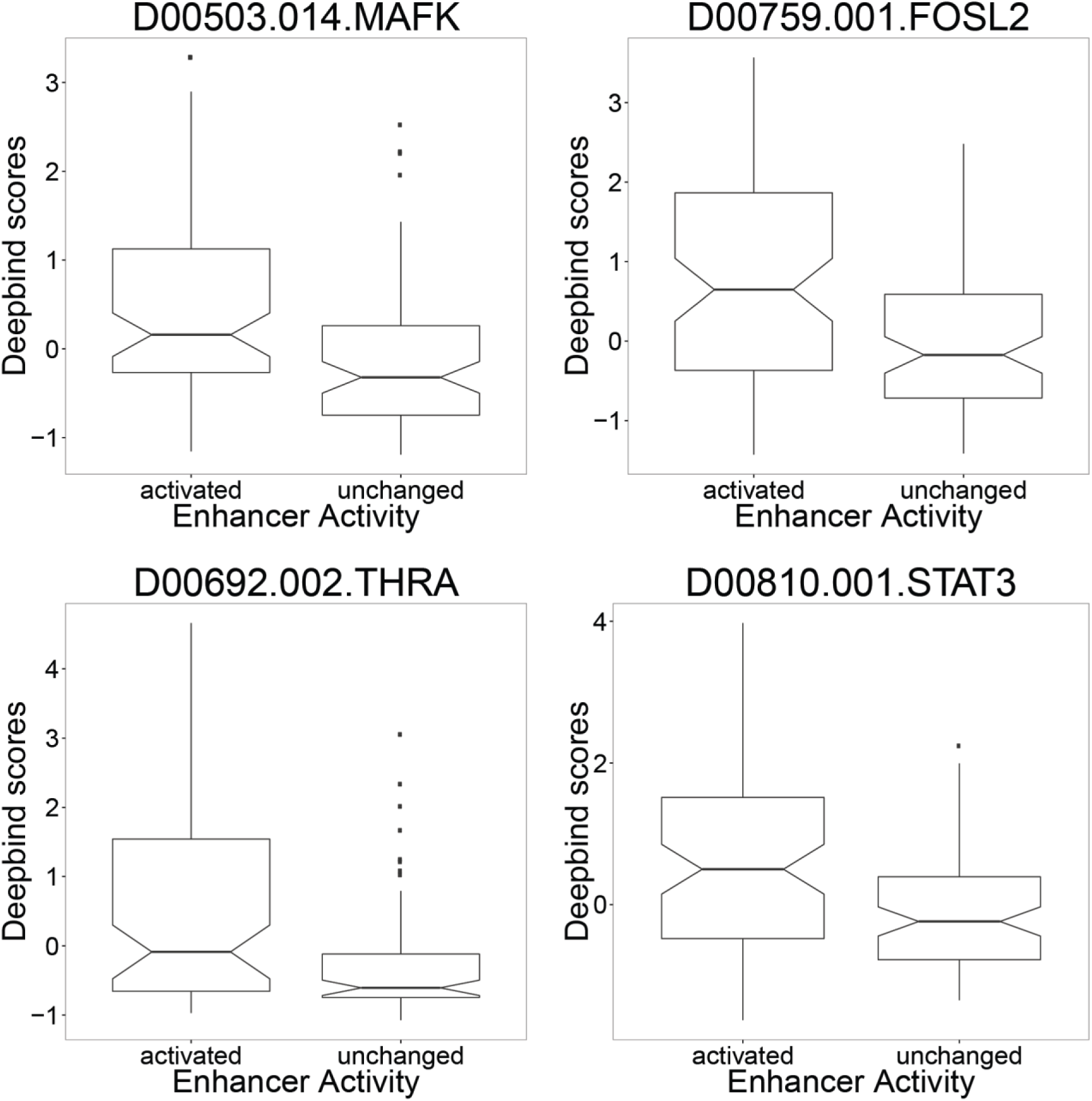
TFs that had statistically different DeepBind scores between enhancers which show activation and those that are not differentially expressed as categorized by levels of post-pause polymerase.

